# Slow-wave sleep and REM sleep differentially contribute to memory representational transformation

**DOI:** 10.1101/2024.08.05.606592

**Authors:** Jing Liu, Danni Chen, Tao Xia, Shengzi Zeng, Gui Xue, Xiaoqing Hu

## Abstract

Memory transforms over time, gradually becoming less idiosyncratic and more gist-like. While sleep contributes to memory transformation, how different sleep stages and EEG activity influence memory transformation is far from clear. Applying representational similarity analysis to electroencephalogram (EEG) recordings, we examined memory representational transformation at both the idiosyncratic “item-level” and the generic “category-level”. Our findings revealed that after an overnight sleep, item-level neural representations for post-sleep remembered items were abolished. In contrast, category-level representations remained prominent, but they became distinctive from pre-sleep. Across participants, more rapid eye movement (REM) sleep relative to slow-wave sleep (SWS) was associated with reduced item-level neural representational strength, increased category-level representational strength, as well as the decreased item-level representational similarity between pre-sleep learning and post-sleep retrieval sessions. Moreover, the theta and beta EEG power during REM sleep, and delta power during SWS differentially supported these representational transformations. These findings suggest that post-learning REM sleep and SWS play differential roles in supporting overnight memory transformation.

## Introduction

Sleep consolidates and transforms newly acquired information, making it long-lasting for future use (Diekelmann & Born, 2010; Rasch & Born, 2013; Stickgold, 2005). While extensive evidence has established the sleep’s benefits in consolidating episodic memory (for meta-analysis see Berres & Erdfelder, 2021), how sleep transforms memory remains unclear. Theoretical models propose that sleep transforms idiosyncratic memory episodes into generalized gist or schema (Dudai et al., 2015; Inostroza & Born, 2013; Landmann et al., 2014; Nadel et al., 2012; Payne, 2011; Stickgold & Walker, 2013; Xue, 2022). This sleep-mediated memory transformation is largely inferred from pre- vs. post-sleep behavioral changes in memory tests implicating integration, generalization, and schematization (Barry et al., 2019; Ellenbogen et al., 2007; Friedrich et al., 2015; Lewis & Durrant, 2011; Payne et al., 2009). Despite these promising findings, behavioral measurements may fall short in characterizing the complexity and fidelity of memory representations (Heinen et al., 2024; Xue, 2022). Therefore, it is desirable to obtain direct neural evidence delineating the sleep-mediate memory representational transformation.

We aim to address this question by leveraging the analytical power of Representational similarity analysis (RSA) to examine memory representations at different levels in the human brain (Diedrichsen & Kriegeskorte, 2017). Specifically, RSA can decompose neural representations of individual items into item- and category-level representations (Lee et al., 2019; Ritchey et al., 2013; Wu & Fuentemilla, 2023). Item-level representations capture neural representations unique to specific stimuli (Kuhl & Chun, 2014), while category-level representations capture neural patterns shared across stimuli within semantic categories (Koutstaal et al., 2001; Naspi et al., 2021). Applying the RSA to EEG recordings, we examined both item- and category-level representations within pre-sleep learning and post-sleep retrieval sessions, respectively (i.e., within-session RSA), and item-/category-level representational similarity between pre- and post-sleep sessions (i.e., cross-session RSA). We hypothesize that sleep facilitates memory transformation and gist-extraction among post-sleep remembered items. Specifically, for within-session RSA, we expected that both item- and category-level representations would be present in the pre-sleep learning session (Liu et al., 2021). Following sleep, we hypothesized that the item-level representation would be diminished and even abolished in the post-sleep retrieval session (i.e., reduced item specificity) (Feld & Born, 2017). In contrast, we anticipated that category-level representation shall persist and remain identifiable in the post-sleep retrieval session (i.e., enduring gist-like information). Moreover, we expected that the cross-session RSA will show lower item-level and/or category-level neural representational similarities as compared to the within-session RSA, as a result of memory representational transformation (Fig. 1A-B).

**Fig 1.**
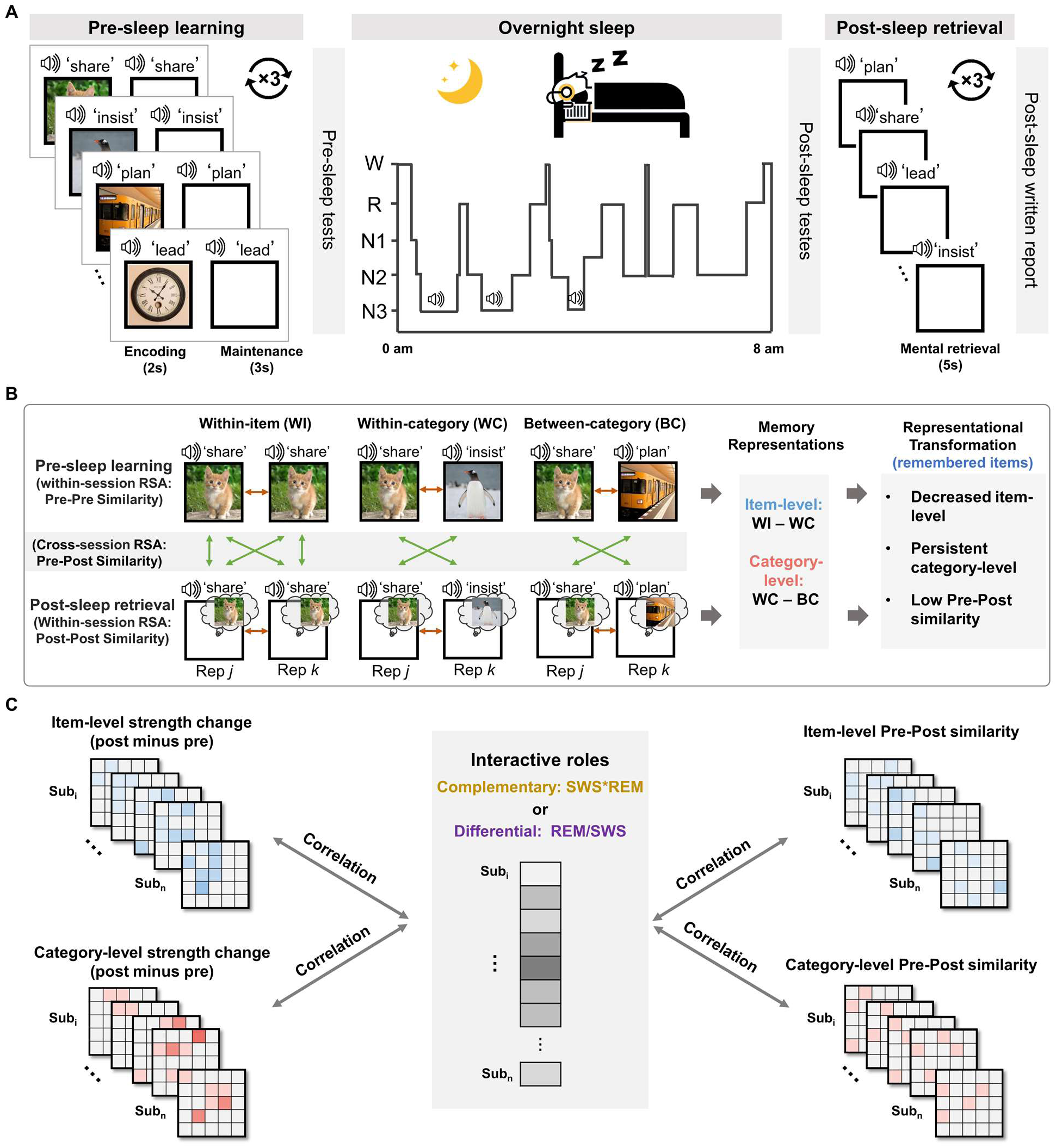
Experimental paradigm and analytic scheme of memory representational transformation across sleep. **(A)** The experimental procedure includes pre-sleep learning (i.e., encoding and maintenance), pre-sleep tests, overnight sleep with TMR cueing during slow-wave sleep, and post-sleep retrieval tests. EEG was recorded throughout the whole experiment. **(B)** Within-session RSA examined memory representations within each of the pre-sleep learning (Pre-Pre similarity) and post-sleep mental retrieval (Post-Post similarity) sessions; Cross-session RSA examined the memory representational similarity between these two sessions (Pre-Post similarity). Both the within-session RSA and cross-session RSA were performed at two different levels: item-level and category-level. Item-level representations were obtained by contrasting the within-item similarity versus within-category similarity (WI-minus-WC), while the category-level representations were obtained by contrasting the within-category similarity versus between-category similarity (WC-minus-BC). We hypothesized that memory representational transformation would be manifested by the following indexes: decreased item-level representations, while persistently prominent category-level representations from pre- to post-sleep; and the low cross-session pre-post similarities. **(C)** If SWS and REM sleep play complementary roles in memory representational transformation, then the product of the REM sleep amount by the SWS amount (SWS*REM) should correlated with memory representational transformation indexes. Otherwise, if REM sleep and SWS play differential roles, then the REM sleep amount relative to the SWS amount (REM/SWS), should be correlated with the memory representational transformation indexes.

More critically, how different sleep stages, particularly the SWS and REM sleep, interactively contribute to memory representational transformation remains controversial. One perspective suggests that these two sleep stages may complement each other in optimizing memory consolidation and transformation (Brodt et al., 2023; Diekelmann & Born, 2010; Giuditta et al., 1995; Inostroza & Born, 2013). Specifically, during SWS, repeated memory reactivation would integrate newly encoded memories into pre-existing memory schema, transforming hippocampal-dependent memory into more neocortex-dependent gist-like representations. Subsequent REM sleep would further stabilize these transformed representations via synaptic consolidation. Supporting this hypothesis, animal studies suggest that SWS-initiated cortical plasticity for memory consolidation is reinforced by the following REM sleep episode (Miyawaki & Diba, 2016; Ribeiro et al., 2007). Moreover, human studies showed that the product of the durations of SWS and REM sleep (SWS * REM), reflecting their complementary roles, explains overnight memory consolidation (Hu et al., 2015; Mednick et al., 2003; Stickgold et al., 2000). If SWS and REM complement each other in memory transformation, then a higher SWS * REM should be associated with greater overnight memory representational transformation (Fig. 1C).

An alternative perspective posits that SWS and REM sleep play differential roles in sleep-mediated memory transformation (MacDonald & Cote, 2021; Payne, 2011). According to this view, SWS stabilizes memory representations in their original formats, while REM sleep primarily refines and transforms them into schema-like formats. Supporting this perspective, research has shown that REM sleep duration is positively associated with schema-conformant memory consolidation and creative problem-solving, while SWS duration showed an opposite trend (Cai et al., 2009; Durrant et al., 2015). Consistent with this idea, a recent study showed that greater memory distortion or modification occurs after REM-rich sleep, while stabilization of the undistorted original memory occurs after SWS-rich sleep (Kaida et al., 2023). If this perspective holds true, a higher REM to SWS duration ratio (i.e., REM/SWS) should be associated with greater memory representational transformation (Fig. 1C).

Here, combining overnight sleep EEG recordings with RSA, we examined item- and category-level memory representational transformation across sleep. Our results revealed substantial memory representational transformation for post-sleep remembered items: while pre-sleep memory representations contained both item-level and category-level content, post-sleep memory representations were predominantly categorical. More importantly, a higher REM/SWS duration ratio was associated with reduced item-level representational strength, increased category-level representational strength, and reduced item-level cross-session similarity. Thus, our findings support the differential roles of SWS and REM in memory representational transformation.

## Results

A total of 35 participants (26 females, mean age ± SD: 22 ± 2.79) were included in the analysis. Participants completed three major task sessions: pre-sleep learning, overnight sleep, and post-sleep mental retrieval tests (see Fig. 1 and Methods). During pre-sleep learning, participants learned 96 unique word-picture pairs, with each repeated three times. After a distraction task, participants were tested for their memory on half of the learning pairs. Subsequently, participants went to nocturnal sleep, during which targeted memory reactivation (TMR) was performed during SWS. In the post-sleep mental retrieval session, participants closed their eyes to mentally recall associated pictures as vividly as possible, prompted by auditory cues. Similar to the pre-sleep learning, each word-picture pair was mentally retrieved three times. Immediately after the mental retrieval task, participants were asked to write down the picture content, promoted by individual printed cue words. Participants showed an average accuracy rate of 0.40 (SD: 0.18) in this task, serving as the post-sleep memory retrieval performance. Note that the TMR cued versus uncued items did not differ in post-sleep retrieval performance (*t*(34) = −1.68; *p* = 0.102). The TMR effect and the associated neurocognitive processing are not the main focus of the study and were reported in Liu et al., 2023.

### Overnight neural representational transformation for post-sleep remembered items

To understand how memory representation transforms across an overnight sleep while remaining retrievable, we first examined the neural representations during the pre-sleep learning session (i.e., including both encoding and maintenance periods) for post-sleep remembered items. Following previous studies (Lee et al., 2019; Liu et al., 2020; Ritchey et al., 2013), we performed the RSA on the auditory cue-elicited EEG power patterns to extract item- and category-level neural representations. These representations capture fine-grained item-specific information and generalized categorical information, respectively. For item-level representations, we contrasted the EEG power pattern similarities between trials of the same pictures (Within-item, WI similarity) versus the similarity between trials of different pictures from the same category (Within-category, WC similarity, see Methods). For category-level representations, we contrasted the WC similarity with the similarity between trials of different pictures from different categories (Between-category, BC similarity) (Fig. 1B).

To examine the temporal dynamics of representational transformation, we computed the similarity values by correlating the EEG power pattern across frequencies (2-40 Hz) and all clean channels between artifact-free learning trials, in 500 ms sliding time windows with a stride of 100 ms during the 5 s post-stimuli epoch. We found significant item-level representations (i.e., WI > WC similarity) within a ∼700-2500 ms cluster and a ∼2000-4000 ms cluster post-stimuli onset (*p*s_cluster_ < 0.032, corrected by the non-parametric cluster-based permutation test, Fig. 2A). We also found significant category-level representations (i.e., WC > BC similarity) within a significant cluster (∼0-1000ms and 3000-4800ms post stimuli onset, *p*_cluster_ = 0.043, Fig. 2B). These results suggested that both item-level and category-level neural representations emerged during pre-sleep learning session for those post-sleep remembered items.

**Fig 2.**
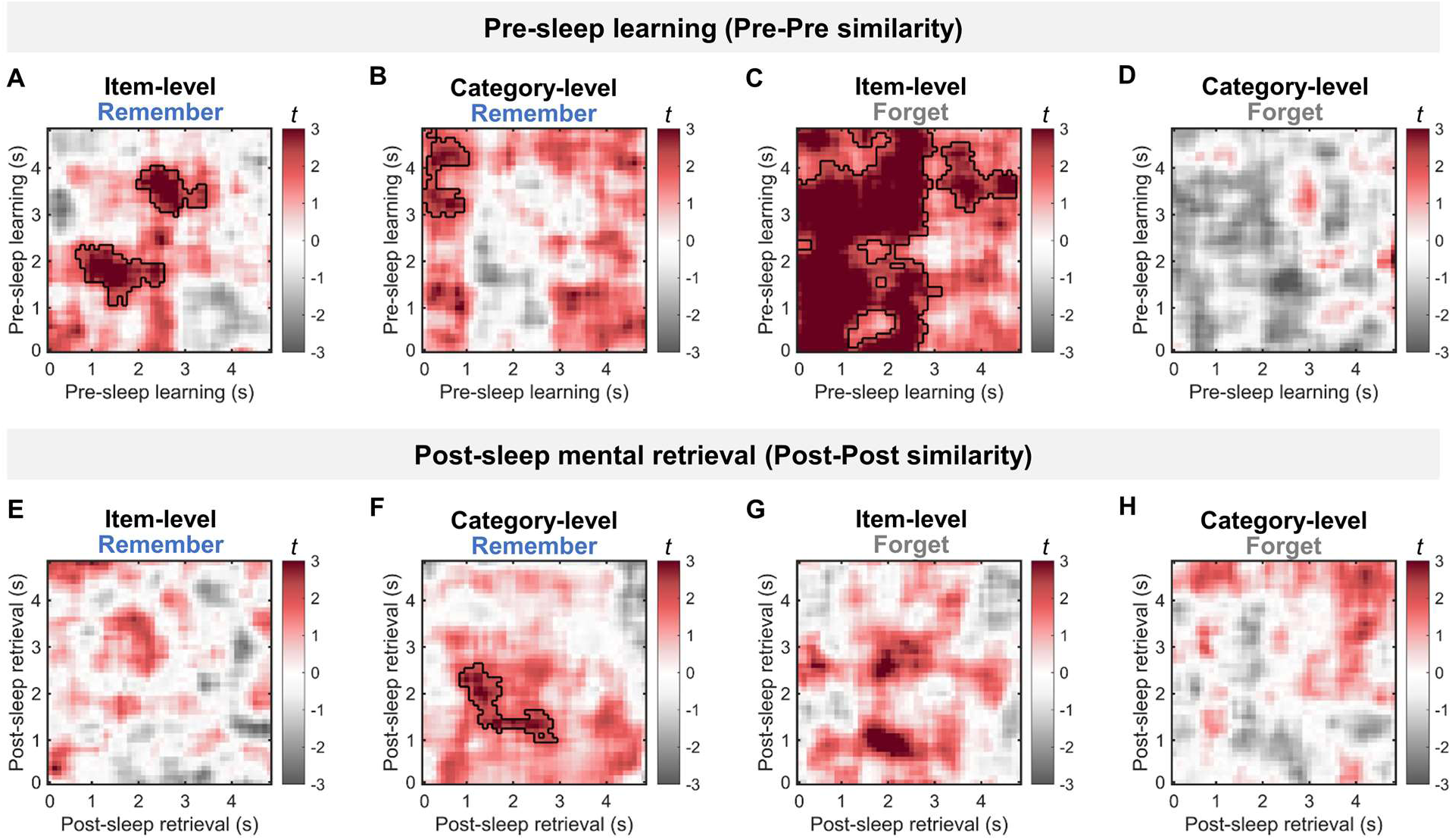
Neural representations within pre-sleep learning and post-sleep mental retrieval sessions, respectively. **(A-B)** During the pre-sleep learning session, significant item-level representations and category-level representations were identified for post-sleep remembered items in the clusters. **(C-D)** For post-sleep forgotten items, item-level but not category-level representations were identified during the pre-sleep learning session. **(E-F)** During the post-sleep mental retrieval session, no item-level representations but significant category-level representations were identified for post-sleep remembered items in the cluster. **(G-H)** For post-sleep forgotten items, neither item-level representations nor category-level representations were identified during the post-sleep mental retrieval session. Significant clusters with *p*_cluster_ < 0.05 were circled by black lines.

As a control analysis, we performed the same RSA for post-sleep forgotten items. The results only revealed two significant clusters showing significant item-level representations (*ps*_cluster_ < 0.042, Fig. 2C), but no significant category-level representations (*ps*_cluster_ > 0.690, Fig. 2D) during pre-sleep learning. Comparisons between post-sleep remembered and forgotten items revealed two clusters showing Remember < Forget item-level representations (all *p*s_cluster_ < 0.050, see Fig. S1) and three clusters showing Remember > Forget category-level representations (all *p*s_cluster_ < 0.027, see Fig. S1), may reflect that greater transforming from item-level to semantic category-level representations during learning predicts better long-term memory (Liu et al., 2021).

Next, we examined item- and category-level neural representations during the post-sleep mental retrieval session. For post-sleep remembered items, contrary to the pre-sleep learning session, we did not find significant clusters showing item-level representations (*p*_cluster_ > 0.455, Fig. 2E). However, we found significant category-level representations during ∼900-2900 ms time window, *p*_cluster_ = 0.045, Fig. 2F). Control analyses on post-sleep forgotten items did not reveal any significant clusters for either item- or category-level representations (all *p*s_cluster_ > 0.064, Fig. 2G-H). Direct comparison between remembered vs. forgotten items only revealed a significant cluster showing Remember > Forget category-level representations (*p*_cluster_ = 0.017), but there was no significant difference for item-level representations (*p*_cluster_ > 0.233, see Fig. S1).

We next performed the cross-session RSA to examine the representational similarity between pre-sleep learning and post-sleep mental retrieval sessions (i.e., Pre-Post Similarity, Fig. 1B). In line while extending previous study (Liu et al., 2021), we did not find any significant Pre-Post representational similarities on either item-level (i.e., WI vs. WC) or category-level (i.e., WC vs. BC) for post-sleep remembered items (all *p*s_cluster_ > 0.487, Fig. 3A). However, post-sleep remembered items showed greater Pre-Post WI and WC similarity than forgotten items (all *p*s_cluster_ < 0.049, see Fig. S2). These results suggested that while successful memory retrieval leads to greater cross-session neural pattern similarity than forgotten items, there were no discernible item-level or category-level representations preserved from pre- to post-sleep session.

**Fig 3.**
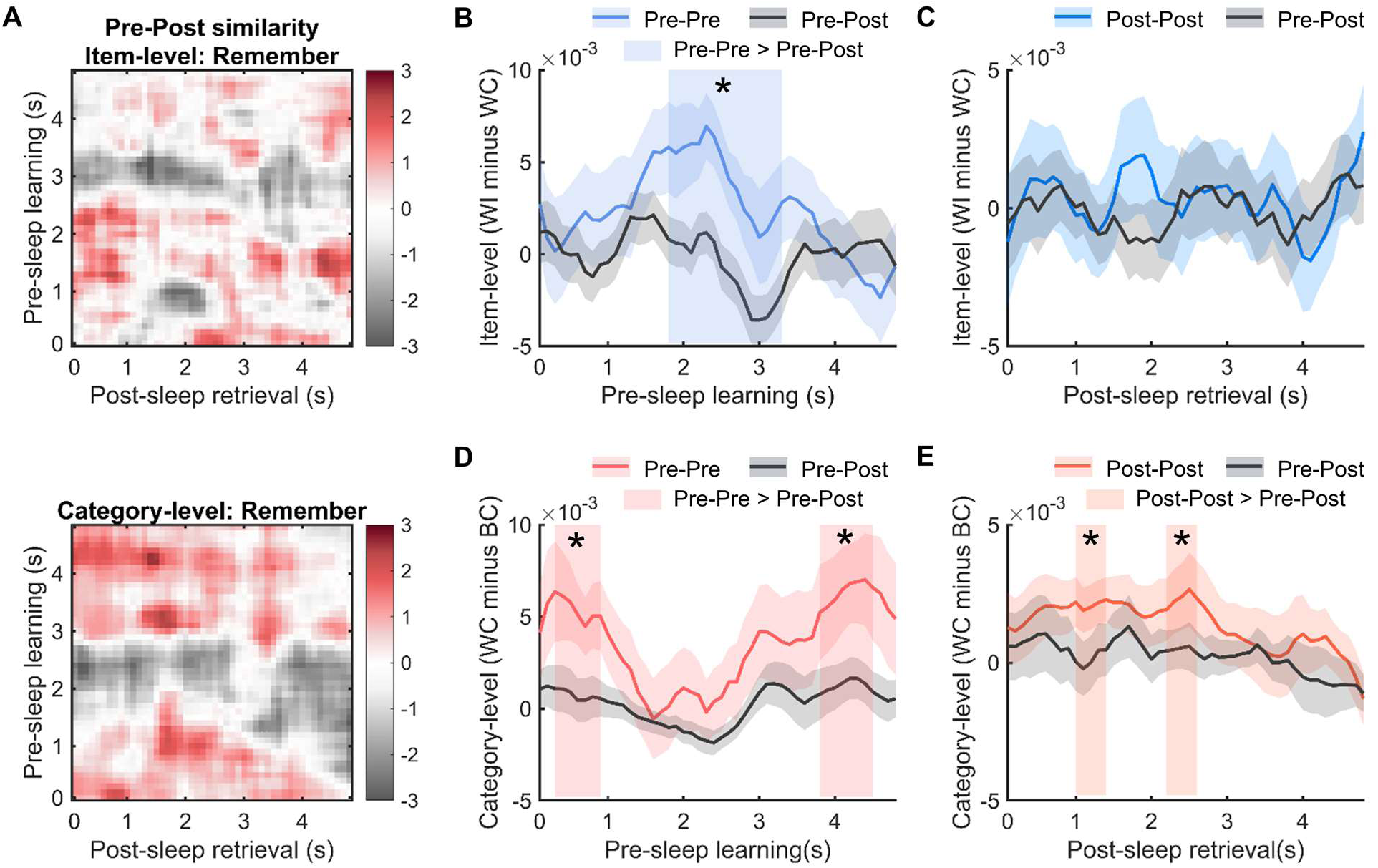
Cross-session representational similarities and their contrast with the within-session representational similarities. **(A)** No significant item-level (upper panel) or category-level (lower panel) Pre-Post similarity for post-sleep remembered items in the cross-session RSA. **(B-C)** The item-level Pre-Post similarity was lower than that within the pre-sleep learning session (i.e., Pre-Pre similarity) but not significantly different from that within the post-sleep retrieval session (i.e., Post-Post similarity). **(D-E)** The category-level Pre-Post similarity was lower than both the category-level Pre-Pre similarity and Post-Post similarity. Significant clusters were indicated by the shaded rectangles. *: *p*_cluster_ < 0.05.

The absence of the cross-session, item-/category-level memory representational similarity could reflect either weakened or changed neural representational patterns across sleep. Previous studies suggested that memory representational pattern changes entailed greater within-session representational similarities than cross-session representational similarities (Liu et al., 2021; Spaak et al., 2017; Stokes, 2015; Xiao et al., 2017). We thus compared within-session similarities (Pre-Pre and Post-Post similarity) with cross-session Pre-Post similarity at the item-level and category-level, respectively. To enable direct comparisons, we averaged the Pre-Pre (see Fig. 2) and Pre-Post similarity (see Figure 3 A) within each pre-sleep learning time window and then contrasted them across pre-sleep learning time windows (see Methods). Similarly, we contrasted Post-Post similarity with Pre-Post similarity across post-sleep retrieval time windows. The results revealed a significant cluster showing greater item-level Pre-Pre similarity than the Pre-Post similarity (*p*_cluster_ = 0.013, Fig. 3B), while no significant difference between item-level Post-Post and Pre-Post similarity (*p* > 0.275, Fig. 3C), which may reflect decayed item-level representations. In addition, for the category-level representations, we found significant clusters indicating that both within-session similarities (i.e., Pre-Pre and Post-Post) were significantly greater than the cross-session Pre-Post representational similarity (Pre-Pre > Pre-Post clusters: *p*s_cluster_ < 0.027; Post-Post > Pre-Post clusters: *p*s_cluster_ < 0.032; Fig. 3D-E, see also Fig. S3). These results suggested that despite significant category-level memory representations within both pre-sleep learning and post-sleep retrieval sessions, they were transformed into distinct formats after an overnight sleep.

To rule out the possibility that pre-sleep testing (on half of word-picture pairs) following pre-sleep learning may influence the overnight memory transformation, we compared the memory representations between pre-sleep tested items and pre-sleep untested items. The results revealed that, among post-sleep remembered items, no significant difference between tested and untested items was found at either the item-level or category-level during the post-sleep mental retrieval session (all *p*s_cluster_ > 0.152, see Fig. S4). In addition, no significant difference was found at either the item-level or category-level Pre-Post similarity (all *p*s_cluster_ > 0.340, see Fig. S4). Similarly, we compared the TMR cued versus uncued items to examine the impact of TMR on memory representations. The results revealed no significant difference at either the item-level or category-level during post-sleep mental retrieval session (all *p*s_cluster_ > 0.196, see Fig. S5) and no significant difference at either the item-level or category-level Pre-Post similarity (all *p*s_cluster_ > 0.195, see Fig. S5).

### REM/SWS, but not SWS*REM, is associated with memory representational transformation for remembered items

We next investigated how SWS and REM sleep influence representational transformation. To answer this question, we first scored the sleep EEG using the toolbox, Yet Another Spindle Algorithm (YASA, Vallat & Walker, 2021), the results of which were further verified by an experienced sleep researcher (see Methods; See Table 1, Fig. S6 for sleep staging). One participant with disconnected EEG recordings during sleep was excluded, resulting in 34 participants in the following data analysis.

We hypothesize that if SWS and REM sleep play complementary roles in memory representational transformation, then the production of SWS% (i.e., percentage in total sleep time) and REM% (SWS*REM) would be associated with memory transformation indexes, i.e., reduced item-level representational strength and the relatively persistent category-level representational strength, and the low Pre-Post similarity for item-level and/or category-level representations. In contrast, if SWS and REM sleep play differential roles in memory transformation, then the REM% relative to the SWS% (i.e., REM/SWS) should be associated with memory representational transformation indexes.

To test these hypotheses, we computed overnight item-level (or category-level) representational strength change by subtracting the mean strength of Pre-Pre item-level (or category-level) representational similarity from Post-Post item-level (or category-level) representational similarity for each participant (i.e., Post minus Pre). We then performed the correlation analysis between SWS*REM and representational strength change across all participants (see Fig. 1C). The results revealed no significant clusters correlating SWS*REM with either item-level strength change or with the category-level strength change (all *p*s_cluster_ > 0.220, corrected by the non-parametric cluster-based permutation test, Fig. 4A-B). In addition, no significant clusters were found correlating SWS*REM with either item-level Pre-Post similarity or category-level Pre-Post similarity (all *p*s_cluster_ > 0.140, Fig. 4C-D).

**Fig 4.**
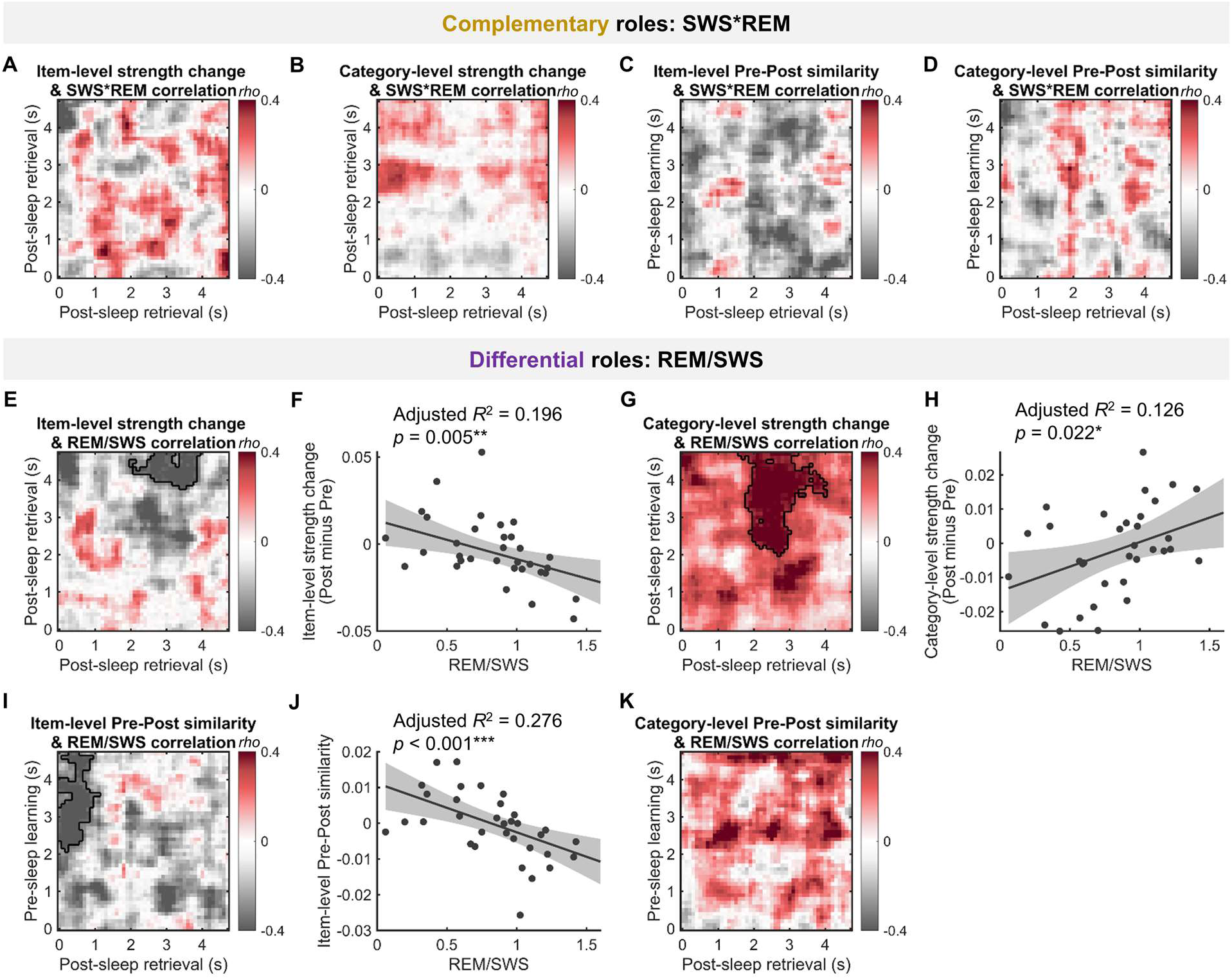
The interactive functional roles of SWS and REM sleep in memory representational transformation. **(A-B)** SWS*REM showed no significant correlation with either item-level or category-level representational strength change (Post minus Pre). **(C-D)** SWS*REM showed no significant correlation with Pre-Post item-level or category-level representation. **(E-F)** REM/SWS was negatively associated with item-level representational strength change. **(G-H)** REM/SWS was positively correlated with category-level representational strength change. **(I-J)** REM/SWS was negatively correlated with item-level Pre-Post similarity. **(K)** No significant clusters were found between category-level Pre-Post similarity and REM/SWS. Significant clusters with *p*_cluster_ < 0.05 were circled by black lines. ***: *p* < 0.001; **: *p* < 0.01; *: *p* < 0.05.

For REM/SWS, we found a significant cluster showing the REM/SWS was negatively correlated with the item-level representational strength change (*p*_cluster_ = 0.028, within the cluster: *β* = −0.022, adjusted *R*^2^ = 0.196, *p* = 0.005, Fig. 4E-F). In contrast, the REM/SWS was significantly positively correlated with the category-level representational strength change (*p*_cluster_ = 0.021, within the cluster: *β* = 0.014, adjusted *R*^2^ = 0.126, *p* = 0.022, Fig. 4G-H). In addition, the results revealed a significant cluster showing that REM/SWS was negatively associated with the item-level Pre-Post cross-sleep similarity (*p*_cluster_ = 0.008, within the cluster: *β* = −0.014, adjusted *R*^2^ = 0.276, *p* < 0.001, Fig. 4I-J), while no significant clusters were observed between REM/SWS and category-level Pre-Post similarity (*p*s_cluster_ > 0.202, Fig. 4K). However, when correlating the memory representational transformation indexes with REM% and with SWS% respectively, we only found a significant cluster showing a negative correlation between item-level Pre-Post similarity and REM% (see Fig. S7), with the explained 25.2% of inter-participant variance lower than the REM/SWS (27.6%). Control analysis between REM/SWS and memory representational transformation indexes among post-sleep forgotten items revealed no significant results (all *p*s_cluster_ > 0.282, see Fig. S8).

These results collectively suggest that SWS and REM sleep play differential roles, instead of complementary roles, in memory representational transformation among post-sleep remembered items. The greater amount of REM sleep, in contrast to SWS, is associated with significant memory representational transformation across participants, as indexed by the reduced item-level representational strength and enhanced category-level representational strength, alongside the reduced item-level cross-session representational similarity.

### REM and SWS EEG power are differentially associated with the neural representational transformation for remembered items

Beyond the REM and SWS duration, we next investigated what electrophysiological activities during REM sleep and SWS modulate this transformation. Prior studies have suggested that during REM sleep, frontal theta and beta activities contribute to memory consolidation (Harrington et al., 2021; Nishida et al., 2009; Vijayan et al., 2017). Building on these results, we calculated the frontal (F3/4 electrodes) theta (4-7 Hz) and beta (15-25 Hz) power relative to the 1-40 Hz total EEG power (see Methods) across all REM sleep epochs. To better examine the effect of REM sleep power across entire overnight sleep on memory representational transformation indexes identified in Fig. 4, we created a comprehensive index of REM sleep power. Specifically, we multiplied the relative power in each frequency band by REM duration, resulting in the total power of each frequency band for REM sleep. The robust linear regression revealed that the total theta power during REM sleep was negatively correlated with item-level representational strength change within the cluster as shown in Fig. 4E (*β* = −29.324, adjusted *R*^2^ = 0.142, *p* = 0.016, Fig. 5A), while positively correlated with category-level representational strength change within the cluster as shown in Fig. 4G (*β* = 21.600, adjusted *R*^2^ = 0.134, *p* = 0.019, Fig. 5B). In addition, the total theta power during REM sleep was negatively correlated with item-level Pre-Post similarity within the cluster as shown in Fig. 4I (*β* = −13.693, adjusted *R*^2^ = 0.109, *p* = 0.032, Fig. 5C). Similarly, total beta band power during REM sleep was negatively correlated with item-level representational strength change (*β* = −197.000, adjusted *R*^2^ = 0.168, *p* = 0.009, Fig. 5D), with a positive but non-significant trend with category-level representational strength change (*β* = 101.700, adjusted *R*^2^ = 0.044, *p* = 0.122, Fig. 5E). In addition, the total beta band power during REM sleep was negatively correlated with item-level Pre-Post similarity (*β* = −130.23, adjusted *R*^2^ = 0.242, *p* = 0.002, Fig. 5F). Note that similar results were found when correlating relative REM theta or beta power with these memory representational transformation indexes (see Fig. S9). Further exploratory analysis with other frequency bands power (i.e., delta: 1-3 Hz; alpha: 8-12 Hz; sigma: 11-16 Hz) during REM sleep and memory representational transformation indexes did not yield significant results (all *p*s > 0.102).

**Fig 5.**
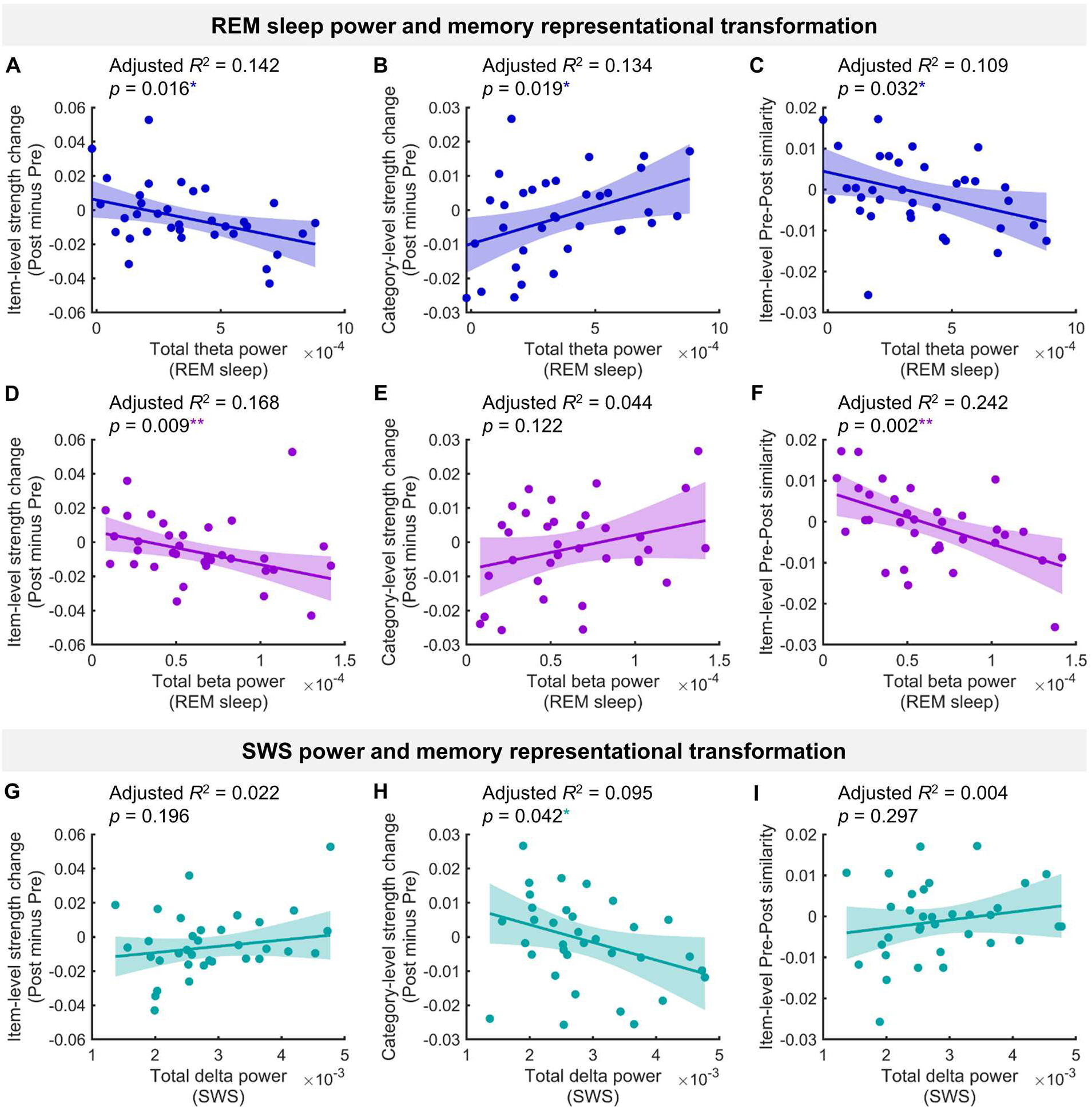
The relationship between sleep EEG power and memory representational transformation. **(A-C)** Overall frontal theta power during REM sleep was negatively associated with the item-level strength change (post minus pre) and positively associated with the category-level strength change (post minus pre) and negatively associated with the item-level Pre-Post similarity. **(D-F)** Overall frontal beta power during REM sleep was negatively associated with the item-level strength change and negatively associated with the item-level Pre-Post similarity. A positive but nonsignificant trend was found between the beta power and category-level strength change. **(G-I)** Overall frontal-central delta power during SWS was negatively correlated with the category-level strength change. No significant correlation was found between delta power and item-level strength change and Pre-Post similarity. **: *p* < 0.01; *: *p* < 0.05.

We next examined, during the SWS, how the canonical frontocentral (Fz and Cz electrodes) 1-4 Hz delta and 11-16 Hz spindle-related sigma power were associated with memory representations transformation indexes. In contrast to REM theta and beta power, total SWS delta power across the overnight SWS sleep (i.e., delta power * SWS amount) was negatively correlated with category-level representational strength change (*β* = −5.181, adjusted *R*^2^ = 0.095, *p* = 0.042, Fig. 5H). No significant correlations were found for item-level representational strength change or for Pre-Post similarities (all *p*s > 0.195, Fig. 5G, I). In addition, no significant correlation was found between total SWS sigma band power and all these memory representational indexes (all *p*s > 0.183). Besides, relative SWS delta power but not sigma power was negatively correlated with category-level representational strength change (see Fig. S10).

## Discussion

Examining neural representations across pre-sleep learning, overnight sleep, and post-sleep retrieval sessions, we demonstrated that memory representations of individual items were substantially transformed. From pre-sleep learning to post-sleep retrieval, idiosyncratic item-level representation was abolished while categorical representations remained prominent. In addition, the Pre-Post cross-session item-level representations were lower than the Pre-Pre item-level representations, and Pre-Post category-level representations were lower than both Pre-Pre and Post-Post category-level representations. Most importantly, we provide compelling evidence that REM and SWS differentially impact memory representational transformation. Specifically, a greater REM sleep to SWS ratio was associated with reduced item-level representational strength, increased category-level representational strength across sleep, and was associated with reduced cross-sleep item-level representational similarity.

First, our study advances the understanding of overnight memory representational transformation, extending prior research on neural representational transformation observed within minutes or a few hours during wakefulness (Cichy et al., 2014; Favila et al., 2018; Liu et al., 2021; Xiao et al., 2017). Previous research has shown that item-level memory representations are evident during both encoding and retrieval sessions that occurred within 1-2 hours of wakefulness (Favila et al., 2018). Our findings extend these studies by showing that, after overnight sleep, item-level memory representations were no longer significant, while category-level memory representations were persistently prominent from pre-sleep learning to post-sleep retrieval sessions. Our results can be well explained by the functional roles of sleep in transforming memory. Specifically, memory reactivation during the sleep-based consolidation process may facilitate the abstract of the gist information (e.g., the concept of animal) from individual items within the same category (e.g., different animal pictures) (Lau et al., 2011; Lewis & Durrant, 2011). These gist-like semantic category representations were then more likely to survive the global synaptic downscaling and to be extracted during the post-sleep wake retrieval (Feld & Born, 2017). Furthermore, we found that cross-session category-level representations were lower than the Pre-Pre and Post-Post category-level representations, suggesting distinct category-level representations between these two sessions (Liu et al., 2021; Spaak et al., 2017; Xue, 2022). These results support the transformative nature of episodic memory over time (Dudai, 2012; Xue, 2022). In addition, previous research has shown that, after overnight sleep, cortical neural pattern similarity between different memory items is enhanced accompanied by increased hippocampal-cortical network connectivity (Cowan et al., 2020). Further study could combine the fMRI and sleep EEG to test whether different brain networks were engaged during sleep-mediated memory representational transformation at the item- and category-level.

Most critically, our study addressed an under-investigated question: how SWS and REM sleep contribute to the sleep-based memory transformation (Diekelmann & Born, 2010; Inostroza & Born, 2013; Landmann et al., 2014; MacDonald & Cote, 2021; Payne, 2011). Despite some studies suggesting the interactive functional role of SWS and REM sleep in memory enhancement (Batterink et al., 2017; Mednick et al., 2003; Stickgold et al., 2000), most sleep research focused on the relationship between a single sleep stage (e.g., either SWS or REM) and behavioral changes, leading to mixed results (Cai et al., 2009; Durrant et al., 2015; Hennies et al., 2017; Ketz et al., 2018; Lau et al., 2010; Pereira et al., 2023; Tamminen et al., 2013). In our study, we systematically examined the complementary and differential roles of SWS and REM sleep in memory representational transformation. Our findings provided novel evidence that a greater amount of REM sleep is associated with greater memory representational transformation, while a reversed pattern was observed for SWS. These results support Payne’s hypothesis (2011) and sleep reinforcement and refinement hypothesis (MacDonald & Cote, 2021), both of which emphasize the critical role of REM sleep in memory representational transformation and the role of SWS in stabilizing memory in its original format. Note that the differential roles of REM sleep and SWS may also vary depending on the tasks, such as the emotional memory task (Cairney et al., 2015), rule abstract task (Pereira & Lewis, 2020), creative problem-solving task (Lewis et al., 2018), as well as being affected by the amount of information being learned pre-sleep (Feld & Born, 2017), which warrants future research.

Regarding the electrophysiological mechanisms, our results showed that both the REM sleep duration and REM sleep theta and beta power contribute to memory representational transformation. Corroborating our findings, previous research showed that both REM sleep duration and REM sleep EEG theta power were positively correlated with the consolidation of schema-conformant memory items (Durrant et al., 2015). In addition, REM sleep amount was positively associated with facilitated semantic processing (Carr & Nielsen, 2015; Stickgold et al., 1999). Together with these studies, our results suggested that REM sleep facilitates memory reorganization within pre-existing semantic networks or schema. Therefore, it results in more gist-like memory representations which may resist interference (Payne, 2011; Tamaki et al., 2020). Our findings also align well with animal studies, which emphasize the REM sleep theta oscillation in memory consolidation. Specifically, the coherence of amygdalocortical theta oscillations during REM sleep predicts the behavioral changes in fear-conditioned cued memory recall (Popa et al., 2010). Selectively suppressing the theta oscillation during REM sleep leads to impaired fear-conditioned contextual memory. In addition to the theta oscillation, a recent human intracranial EEG study has observed beta oscillations during REM sleep (Vijayan et al., 2017), which couple with theta activity (Cox et al., 2019). In line with these studies, our findings showed similar functional roles of REM sleep theta and beta activities in memory representational transformation.

Notably, sleep-mediated memory representational transformation was specifically documented among post-sleep remembered items, whereas no such effect was found among post-sleep forgotten items. Thus, the documented transformation is an adaptive process supporting long-term memory (Xue, 2022). Previous studies suggested that SWS duration as well as the sigma and delta power during SWS were associated with overall better memory retention, as indicated by behavioral performance changes across all post-sleep remembered and forgotten items (Holz et al., 2012; Scullin, 2013). Although we focused on memory representations rather than overall memory performance, our supplementary analysis did reveal individual differences in the duration of SWS were positively correlated with better memory retention across all pre-sleep tested items (see Fig. S11). However, among post-sleep remembered items, we found the greater delta power during the SWS was associated with lower post-sleep category-level representational strength compared to pre-sleep. While the active systems consolidation model proposes that memory representations are repeatedly reactivated during SWS, possibly facilitating the memory transformation into more gist-like representations (Born & Wilhelm, 2012), our study suggests that this SWS-mediated process alone may not necessarily facilitate the gist-like representational formation.

While participants were capable of retrieving the individual pictures associated with their corresponding auditory cues in the post-sleep written report, we did not find significant item-level neural representations during post-sleep mental retrieval for remembered items. Diminished item-level memory representations could be due to global synaptic down-scaling during sleep (Tononi & Cirelli, 2014). As a result, these neural representations may be less likely to be detected using scalp EEG. In addition, the current study included TMR during the SWS (Liu et al., 2023). However, the functional roles of SWS and REM sleep in modulating memory representational transformation are unlikely to be driven by TMR, given the non-significant cued vs. uncued differences at item-/category-level representations from both the Post-Post and the Pre-Post RSA (see Fig. S5). Nevertheless, we acknowledge that our study cannot rule out the possibility that TMR that occurs during SWS may trigger memory representations into labile states which allows memory representations to be transformed during subsequent REM sleep (Batterink et al., 2017; Tamminen et al., 2017). Future studies should further examine the interactive functional roles of REM and SWS in memory representational transformation during spontaneous overnight sleep.

Overall, our study demonstrates overnight memory transformation: while memory representations containing both item- and category-level representations during pre-sleep learning, only category-level representations were dominant post-sleep. More importantly, REM sleep and SWS played differential roles in the representational transformation: the greater amount of REM sleep, relative to the SWS, was associated with greater memory representational transformation. These findings advance our understanding of the interactive functional roles of human SWS and REM sleep in memory consolidation and transformation.

## Methods

### Participants

Thirty-five healthy, right-handed participants were included in the study (females, mean age ± SD: 22 ± 2.79 years). Two additional participants who exhibited significant body movements during pre-sleep learning/ post-sleep mental retrieval sessions were excluded during initial data visual inspection and screening. Behavioral and wakefulness EEG data analysis were performed on all the 35 included participants. However, for the sleep EEG data analysis, one participant was excluded due to disconnected EEG recordings in the middle of overnight sleep, resulting in a final sample size of 34 participants in sleep analyses. Prior to participation, all participants underwent pre-screening for sleep quality using the Pittsburgh Sleep Quality Index (PSQI) and the Insomnia Severity Index (ISI), ensuring overall good sleep quality. They had not taken any sleep-aid medicines in the past month prior to the experiment. All participants were not diagnosed with any neurological or psychiatric disorders and had normal or corrected-to-normal vision. The study was approved by the Research Ethics Committee of the University of Hong Kong. All participants gave written informed consent prior to participation.

### Experimental design

The experiment encompassed three primary sessions: (1) a pre-sleep session, including a word-picture associative learning task and pre-sleep memory tests, (2) an overnight sleep session with targeted memory reactivation (TMR) administered during NREM sleep for the initial 3-4 sleeping hours, and (3) a post-sleep session, including a post-sleep mental retrieval task and memory tests. All the behavioral tasks were administered using PsychoPy (version: 2020.2.10; https://www.psychopy.org/).

During the pre-sleep word-picture associative learning task, participants were instructed to memorize a total of 96 distinct word-picture pairs. The 96 words were two-character Chinese verbs, while the corresponding pictures were naturalistic images. Each picture fell into one of four categories, namely animals, electronic devices, plants, and transportation tools, with 24 pictures in each category. Each word was randomly paired with a picture for each participant. Each learning trial consisted of three phases: encoding, maintenance, and vividness rating. During encoding, participants were presented with a fixation cross for 0.3 s, followed by a black screen with jittering durations between 0.9 to 1.5 s. Subsequently, a picture was displayed at the center of the screen for 2 s, accompanied by the auditory presentation of the corresponding spoken verb. Participants were explicitly instructed to focus their attention on the picture and memorize the associations between the verbs and the pictures. In the immediately following maintenance period, the presented picture disappeared, and participants were instructed to vividly mentally maintain the picture for a duration of 3 s while hearing the spoken verb again. In the vividness rating phase, participants were required to evaluate the subjective vividness of the mental image they held during the maintenance period on a scale from 1 (not vivid at all) to 4 (very vivid) within 2 s. The entire pre-sleep learning task consisted of three blocks, with each block consisting of 32 distinct verb-picture pairs and each pair repeated three times within a block. To minimize the potential influence of the recency effect, participants engaged in a ∼5-minute math task immediately after completing the learning task.

After the distractor math task and a short break (∼5 minutes), half of the pairs (i.e., 48 pairs) were tested via the cued category-report task and the cued recognition task pre-sleep. In the cued category-report task, each trial started with a 0.3 s fixation, followed by a blank screen (0.9-1.5 s). The spoken verb was played, prompting participants to report whether they “remember” or “forget” the corresponding picture. This stage was self-paced so that participants had enough time to recall. Following the “remember” or “forget” response, participants were asked to report the category of the picture by pressing one of four buttons, with each button representing one of the four categories. In the cued recognition task, the same half of the pairs were tested. Each recognition trial began with a fixation (0.3 s) and was followed by a blank screen (0.9-1.5 s). Participants were then presented with the picture (either a pre-learned picture or a similar lure picture) in the center of the screen while simultaneously hearing the corresponding spoken verb. They were asked to indicate if the picture was the same picture paired with the verb during the previous learning task by pressing the “Yes” or “No” button. After the pre-sleep test, participants went to sleep from approximately 12 am to 8 am. Targeted memory reactivation cueing was delivered during SWS in the first 3-4 hours after participants fell asleep (see Liu et al., 2023 for more details).

Approximately 30 minutes after awakening in next morning, participants were tested on all 96 pairs. The post-sleep test included the same cued recall and cued recognition tasks, with an additional mental retrieval task in between. The mental retrieval task was particularly designed to examine the neural representations during the post-sleep retrieval task. Specifically, participants were asked to keep their eyes closed throughout the entire testing block, during which they were asked to mentally retrieve the associated picture as vividly as possible while hearing the auditory verbs, without any explicit behavioral responses. These auditory verbs were randomly played via the speaker with an interstimulus interval (ISI) of 5 ± 0.2 s, comparable to the trial length during pre-sleep learning. Each auditory cue was repeated three times. After the completion of the mental retrieval task, participants were provided with a printed form containing all the cue verbs presented during the mental retrieval task. They were then asked to write down the specific content they retrieved during the mental retrieval task for each cue verb. We then code the participants’ retrieval performance as follows: if the written content accurately represented the central elements of the corresponding picture, it was labeled as “remember”; if the written content described was incorrect or left blank, it was labeled as “forget”. TMR cued and uncued items (*t*(34) = −1.68; *p* = 0.102) showed no significant difference in the post-sleep retrieval performance, we thereby performed all the analyses after combining cued and uncued items in individual participants.

### EEG recording and preprocessing

EEG data were continuously collected from pre-sleep learning to post-sleep tests, including the overnight sleep, using the amplifier from the eego system (ANT neuro, Netherlands, https://www.ant-neuro.com). Data were sampled at 500 Hz using 64-channel waveguard EEG caps, among which 61 channels were mounted in the international 10-20 system, while two electrodes were placed on the left and right mastoids, and one electrode was positioned above the left eye for EOG measurements. During the sleep EEG recordings, two additional electrodes were placed on both sides of the chin to measure the electromyogram (EMG) using a bipolar reference configuration. Prior to EEG recordings, impedance levels for all channels were maintained below 20 KΩ. During online EEG recordings, the default reference channel (CPz) was used. Offline preprocessing of the EEG data was conducted using the EEGLAB (https://sccn.ucsd.edu/eeglab/) and Fieldtrip (https://www.fieldtriptoolbox.org/) toolboxes, as well as custom MATLAB code.

Specifically, EEG data were first notch filtered at 50 ± 2 Hz, and then bandpass filtered between 0.5 and 40 Hz. The continuous EEG data during the pre-sleep learning task were segmented into epochs spanning from 3000 ms before until 8000 ms after stimulus onset. This long epoch was used to eliminate the edge effect in the subsequent time-frequency analysis. Our main interesting time windows for the pre-sleep learning data are from 0 to 5 seconds relative to the stimulus onset. Similarly, for the post-sleep mental retrieval data, the continuous EEG data were segmented into epochs spanning from 3000ms before until 8000ms after the auditory word onset, with our interesting time windows from 0 to 5000ms post auditory word onset. Epochs affected by the muscle movements were visually inspected and excluded from further analysis. Eye blinks and movements were corrected using the independent component analysis. Any identified bad channels were interpolated using spherical interpolation in EEGLAB. Subsequently, the EEG data were re-referenced to the average of the artifact-free data across all channels. For both the pre-sleep learning and the post-sleep retrieval data, EEG epochs were categorized into post-sleep remembered or forgotten trials based on the accuracy of the written reporting immediately following the post-sleep mental retrieval.

### Sleep scoring

Sleep scoring was conducted on non-overlapping 30-second epochs using the Yet Another Spindle Algorithm (YASA), an open-source, machine learning-based toolbox known for its high performance in sleep analysis (Vallat & Walker, 2021). Prior to sleep scoring, bad channels in the EEG data were marked and interpolated. To align with the recommendations of the YASA toolbox, EEG data were re-referenced to FPz. For sleep scoring, the C4 electrode, as well as the EOG and EMG channels, were used as inputs to the YASA algorithm. The scoring results were then double-checked and corrected by an experienced sleep researcher to ensure accuracy and reliability.

### Time-frequency analysis

For both the pre-sleep learning and post-sleep mental retrieval stages, the epoched EEG data underwent time-frequency analysis using complex Morlet wavelets (six cycles). The frequency range of interest was from 2 to 40 Hz, with a step size of 1 Hz. The time window of interest was from −1 to 5 seconds relative to the stimulus onset. To obtain the spectral power, the magnitudes of the complex wavelet transform were squared. The power data were then normalized by subtracting the mean power in the baseline time windows and then dividing by its mean power within each frequency bin and each channel. For both the pre-sleep learning and post-sleep mental retrieval EEG data, the baseline time window was defined as −0.7 to −0.4 seconds relative to the stimulus onset. All spectral power data were subsequently downsampled to 100 Hz and re-segmented into 5-second epochs, specifically [0 to 5 s] relative to the stimulus onset. The spectral power within this broad frequency range [2 to 40 Hz] and epoch duration were used as features for subsequent representational similarity analyses.

### Oscillatory power estimation during SWS and REM sleep

For sleep EEG data, we first epoched the continuous sleep EEG into 30-second epochs. To separate the oscillatory power from the 1/f power-law effect (i.e., fractal component), we employed Irregular-Resampling Auto-Spectral Analysis (IRASA, Wen & Liu, 2016)). Specifically, for each raw sleep epoch data, IRASA first segmented them into ten equally sized, partially overlapped segments, with each covering 90% of the epoch. It then computed the power spectral density (PSD) of these segments of the raw data using the fast Fourier transform (FFT) with the function of a Hanning window. Afterward, it irregularly resamples each segment by factors of h (ranging from 1.1 to 1.9 in increments of 0.05) and 1/h. It uses cubic spline interpolation for irregular upsampling and anti-aliasing low-pass filtering followed by cubic spline interpolation for irregular downsampling. Then the PSD of the resampled data was computed using the same FFT. It then calculated the geometric mean of the auto-power spectra for each h value across upsampled and downsampled signals for each segment. The median of the power spectral with all h-values for each frequency was obtained to estimate the power spectrum of the 1/f power-law effect (i.e., fractal component). We then average the estimated power spectrum of the fractal component and the original signal’s power spectrum across all time segments for each sleep EEG epoch. The oscillatory power of the PSD for individual epochs was then estimated by subtracting the average power spectrum of the fractal component from the PSD of the raw data. Oscillatory power between 1 to 40 Hz for SWS and for REM sleep was obtained by averaging the oscillatory component across epochs labeled as ‘N3’ (SWS) and ‘REM’, respectively. Canonical sleep oscillations were defined as follows: delta band (1-3 Hz), theta band (4-7 Hz), sigma band (11-16 Hz), and beta band (15-25 Hz). For REM sleep, frontal theta and beta band power were calculated by averaging the oscillatory power from F3 and F4 channels (Marquis et al., 2017; Nishida et al., 2009; ten Brink et al., 2023). For SWS, delta and sigma band power were calculated by averaging the oscillatory power from Fz and Cz channels (Mander et al., 2015; Marshall et al., 2003).

### Representational similarity analysis (RSA)

To analyze the neural representations over time, the RSA was performed between the artifact-free trials by correlating the spectral power across frequencies (i.e., 2-40 Hz) and across all scalp channels in sliding time windows. For both the pre-sleep learning and post-sleep retrieval sessions, the length of sliding time windows was 500 ms, with an incremental step size of 100 ms. To increase the signal-to-noise ratio, the spectral power was averaged across time points within each time window, as in previous studies (Liu et al., 2021). This resulted in a set of features consisting of 39 (frequency) by 61 (channel) values for each time window. Then, for each time window, we calculated the similarity between vectorized features of every two trials that were using Spearman’s corrections. All the correlation values were Fisher *Z*-transformed before further statistical analysis.

We categorized representational similarity values into three types: within-item (WI) similarity, within-category (WC) similarity, and between-category (BC) similarity. These categories were based on the corresponding pictures used in the trial pairs for calculating similarity values. WI similarity refers to the similarity between two trials that share the same pictures. WC similarity refers to the similarity between two trials that involve different pictures from the same category. BC similarity refers to the similarity between two trials that involve pictures from different categories. Comparing WI similarity to WC similarity enables us to examine the item-level neural representations while comparing WC similarity to BC similarity enables us to examine the category-level representations (Lee et al., 2019; Ritchey et al., 2013).

The representational similarity was computed either across repetitions within each task session (i.e., within-session RSA) or between two different sessions (i.e., cross-session RSA). Within the pre-sleep learning session, RSA was conducted on distinct learning trials, yielding the Pre-Pre similarity (see Fig. 1). Within the post-sleep mental retrieval session, the RSA was performed between different post-sleep mental retrieval trials, resulting in the Post-Post similarity. Cross-session RSA was conducted between trial pairs, with one trial originating from the pre-sleep learning session and the other from the post-sleep retrieval session. This analysis yielded the Pre-Post similarity. Given that both within-session RSA and cross-session RSA allowed us to compute WI, WC, and BC similarities, we could examine item-level and category-level representations for Pre-Pre similarity, Post-Post similarity and Pre-Post similarity.

To enable the comparisons of the within-session representational similarity with cross-session representational similarity, we compared the Pre-Pre similarity versus Pre-Post similarity across pre-sleep learning time windows and compared the Post-Post similarity versus Pre-Post similarity across post-sleep mental retrieval time windows. Specifically, for a Pre-Pre two-dimensional similarity matrix *C* (either item-level or category-level representations, see Fig. 2), we first averaged the matrix along its first dimension *j*, resulting in the averaged similarity values for each pre-sleep learning time window in the second dimension *i*:

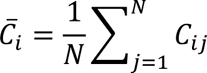

Similarly, we averaged the similarity matrix *C* along the second dimension *i*, resulting in the averaged similarity values in the first dimension *j*:

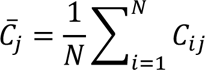

We then obtained the Pre-Pre similarity values for individual pre-sleep learning time windows by further averaging the 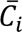 and 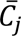. For the Post-Post similarity, we used the same computation to obtain the item-level and category-level similarity values for individual post-sleep retrieval time windows. For the Pre-Post similarity matrix, we averaged it within each pre-sleep learning time window, enabling us to compare it with the Pre-Pre similarity across pre-sleep learning time windows. We averaged the Pre-Post similarity matrix within each post-sleep retrieval time window, enabling us to compare it with the Post-Post similarity across post-sleep retrieval time windows.

For the statistical analysis of representational similarity values across consecutive time windows, multiple comparison corrections were applied using cluster-based nonparametric tests in MATLAB (Maris & Oostenveld, 2007). Specifically, we first conducted statistical tests, such as paired *t*-tests, between conditions (e.g., WI vs. WC or WC vs. BC) within individual time windows. Adjacent time points with significant statistical values (*p* < 0.05) were grouped to form clusters, and cluster-level statistics were calculated by summing t-values within clusters. To determine cluster significance, a null distribution of cluster-level statistics was generated by randomly permuting condition labels 1000 times. For each permutation, the maximum cluster-level statistic was identified. In cases where no significant cluster was observed in a permutation, a value of 0 was assigned. The proportion of cluster-level statistics in the null distribution exceeding the empirical cluster-level statistic determined nonparametric significance.

### Correlation analysis between REM/SWS (or SWS*REM) and representational transformation indexes

We obtained the REM/SWS by dividing the REM amount (i.e., the percentage of REM sleep in total sleep time) by the SWS amount (i.e., the percentage of SWS in total sleep time). Similarly, the SWS*REM was obtained by multiplying the SWS amount by the REM sleep amount. There are four pre-defined representational transformation indexes in the study: item-level representational strength change (post minus pre), category-level representational strength change (post minus pre), item-level Pre-Post similarity, and category-level Pre-Post similarity. To calculate item-level and category-level representational strength changes, we first obtained averaged Pre-Pre similarity matrices across trial pairs for each participant during pre-sleep learning, separately for item-level and category-level representations. Each similarity matrix was then further averaged across all 5-second time windows, resulting in a mean value representing either the pre-sleep item-level or the category-level representational strength for each participant. For each participant, we also obtained averaged item-level and category-level Post-Post similarity matrices across trial pairs. The mean strength of the pre-sleep item-level or category-level representational similarity was then subtracted from the corresponding Post-Post similarity matrices, resulting in item-level and category-level representational strength change matrices. For each time window, we computed Spearman’s correlations between representational strength changes and REM/SWS (or SWS*REM) across participants (see Fig. 1C). Similarly, for each window of the Pre-Post similarity matrices, we performed the Spearman’s correlation between the item-level (or category-level) similarity values with the REM/SWS (or SWS*REM) across participants. These analyses resulted in correlation coefficient matrices, which were then Fisher *Z*-transformed. The correlation analysis across time windows was corrected for using cluster-based nonparametric tests as mentioned above. Briefly, the empirical cluster-level statistics were obtained by summing the transformed correlation coefficients across adjacent time windows with significant correlation (i.e., *p* < 0.05). The null distribution of cluster-level statistics was obtained by shuffling the order of participants for REM/SWS (or SWS*REM) 1000 times while keeping the order of participants for the representational transformation index matrix unchanged. For each shuffling, the same correlation analysis was conducted, and the maximum cluster-level statistic was identified. The nonparametric significance of a cluster was determined by calculating the proportion of cluster-level statistics in the null distribution exceeding the empirical cluster-level statistic.

## Acknowledgment

The research was supported by the National Natural Science Foundation of China (No. 32330039) to G.X.; and Ministry of Science and Technology of China STI2030-Major Projects (No. 2022ZD0214100), National Natural Science Foundation of China (No. 32171056), and General Research Fund (No. 17614922) of Hong Kong Research Grants Council, X.H.; and GuangDong Basic and Applied Basic Research Foundation (No. 2024A1515012667) and Start-up Fund for RAPs under the Strategic Hiring Scheme at The Hong Kong Polytechnic University (Project ID: P0043338) to J.L.

## Author contributions

Conceptualization, J.L. and X.H.; methodology, J.L.; investigation, J.L.; formal analysis, J.L.; writing – original draft, J.L. and X.H.; writing – review & editing, J.L., D.C., T.X., S.Z., G.X. and X.H.; resources, J.L. and X.H.; visualization, J.L. and D.C.; supervision, X.H.; funding acquisition, J.L., G.X., and X.H.

