## Supplementary Materials for "Slow-wave sleep and REM sleep differentially contribute to memory representational transformation"

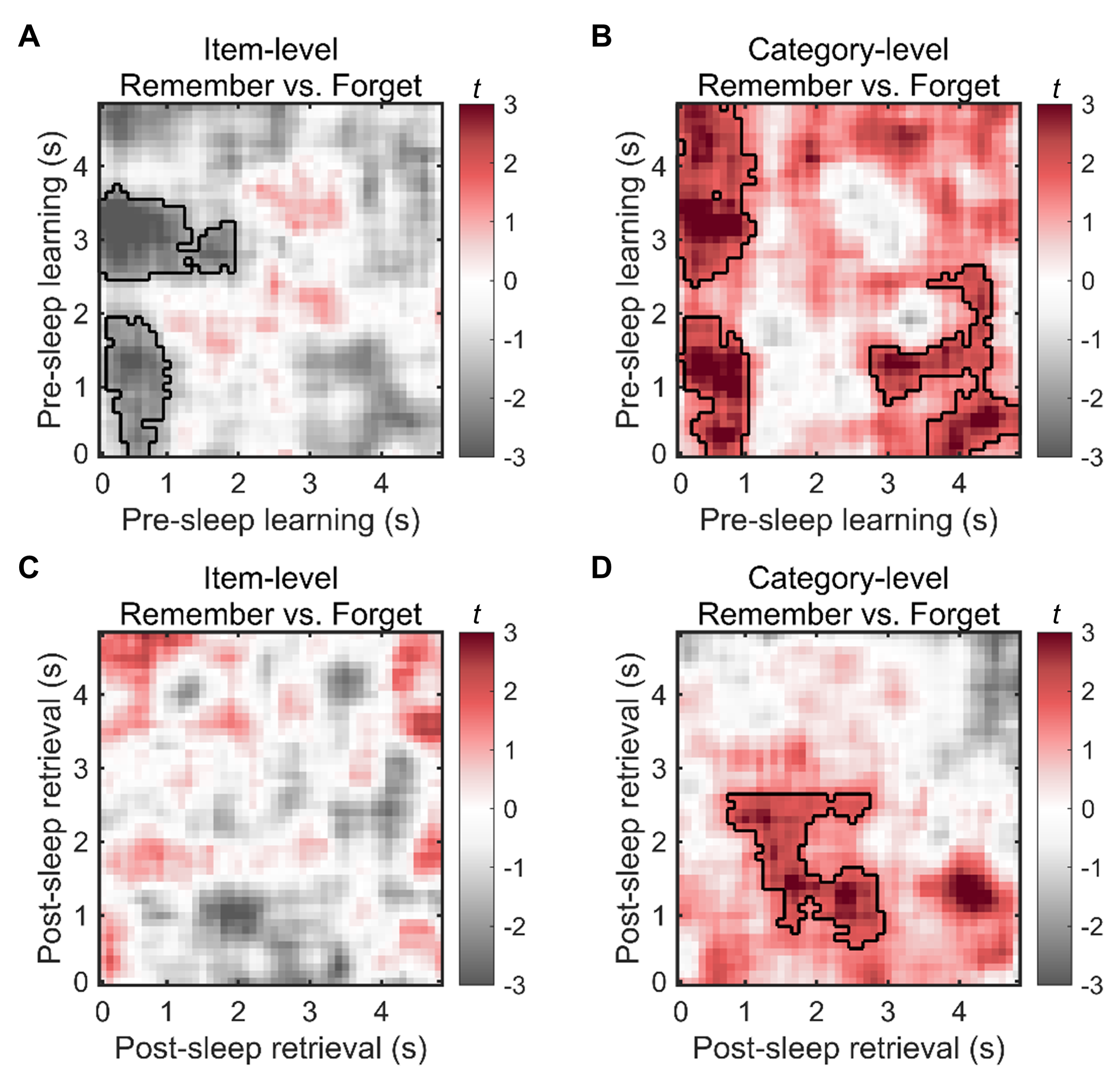

**Fig S1. Item-level and category-level representational differences between post-sleep remembered versus forgotten items. (A-B)** During the pre-sleep learning session, post-sleep remembered items showed lower item-level representations (all *p*s_cluster_ < 0.050) while greater category-level representations (all *p*s_cluster_ < 0.027), as compared to post-sleep forgotten items. **(C)** During the post-sleep mental retrieval session, item-level representations were not different between post-sleep remembered and forgotten items (*p*_cluster_ > 0.233). **(D)** During the post-sleep mental retrieval session, category-level representations were greater for post-sleep remembered items than for forgotten items (*p*_cluster_ = 0.017). Significant clusters with *p*_cluster_ < 0.05 were circled by black lines.

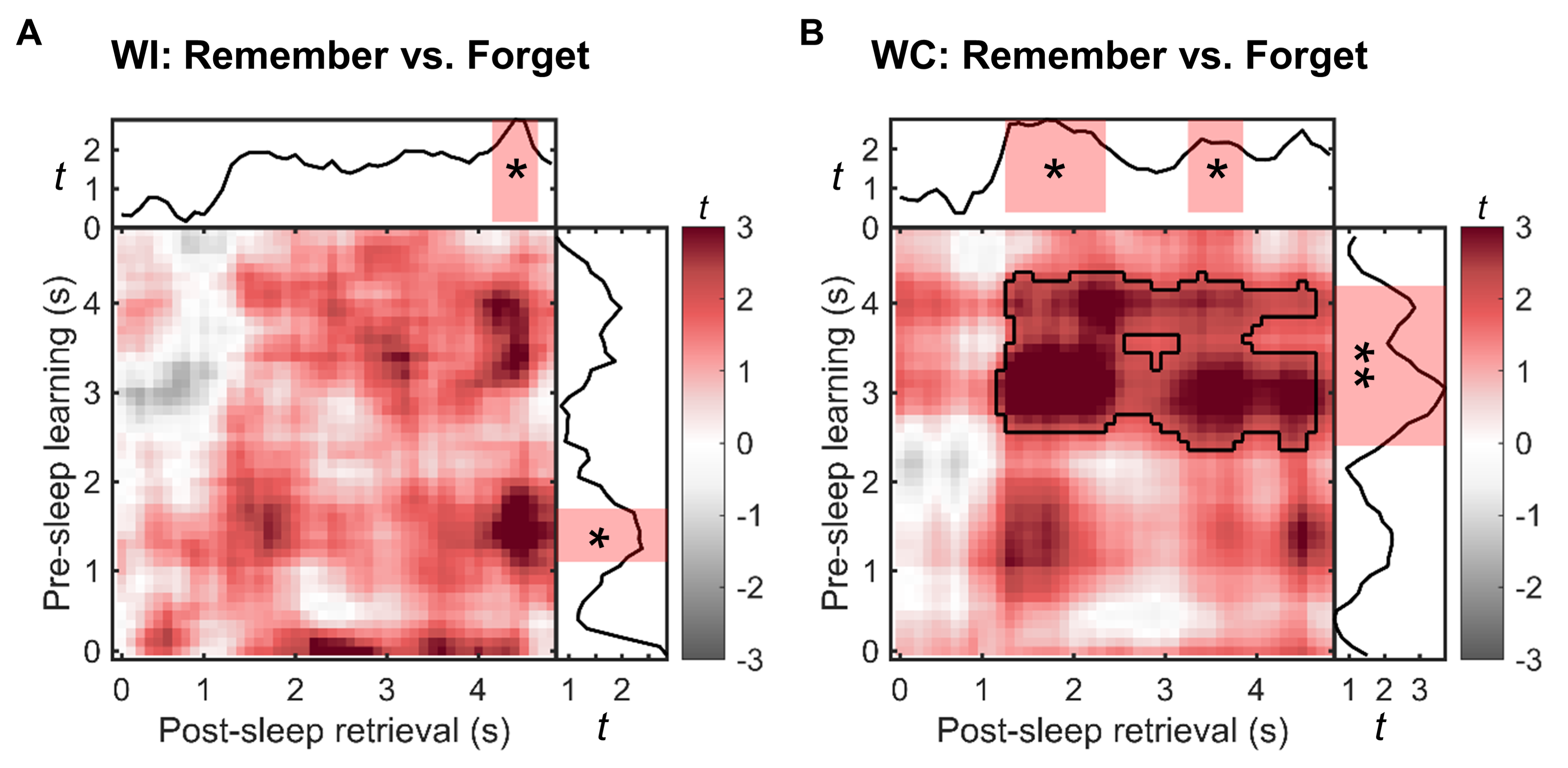

**Fig S2. Greater neural pattern similarity between pre-sleep learning and post-sleep retrieval sessions for remembered than forgotten items**. **(A)** A two-dimensional cluster-based permutation test contrasting the within-item (WI) Pre-Post similarity for remembered versus forgotten items revealed no significant clusters (*p*_cluster_ > 0.110). To enhance the signal-to-noise ratio, we first averaged the WI Pre-Post similarity across all post-sleep retrieval time windows and then contrasted WI similarity between remembered and forgotten items within individual pre-sleep learning time windows. This analysis revealed a significant cluster showing greater WI similarity for remembered than forgotten items in the 1200-1700 ms post-stimulus onset during the pre-sleep learning (*p*_cluster_ = 0.039, indicated by the shaded red rectangle). In addition, after averaging the Pre-Post similarity across all pre-sleep learning time windows, we found greater WI similarity for remembered than forgotten items in the 4200-4700 ms post-stimulus onset during the post-sleep mental retrieval session (*p*_cluster_ = 0.048, indicated by the shaded red rectangle). **(B)** A two-dimensional cluster-based permutation test revealed a significant cluster showing greater within-category (WC) Pre-Post similarity for remembered than forgotten items (*p*_cluster_ = 0.006). After averaging the WC Pre-Post similarity across all post-sleep retrieval time windows, we found a significant cluster showing greater WC similarity for remembered than forgotten items in the 2500-4200 ms post-stimulus onset during the pre-sleep learning session (*p*_cluster_ = 0.006, indicated by the shaded red rectangle). Furthermore, after averaging the WC Pre-Post similarity across all pre-sleep learning time windows, we observed greater WC similarity for remembered than forgotten items in the 1300-2300 ms (*p*_cluster_ = 0.015, indicated by the shaded red rectangle) and 3300-3800 ms (*p*_cluster_ = 0.042, indicated by the shaded red rectangle) post-stimulus onset during the post-sleep retrieval session.

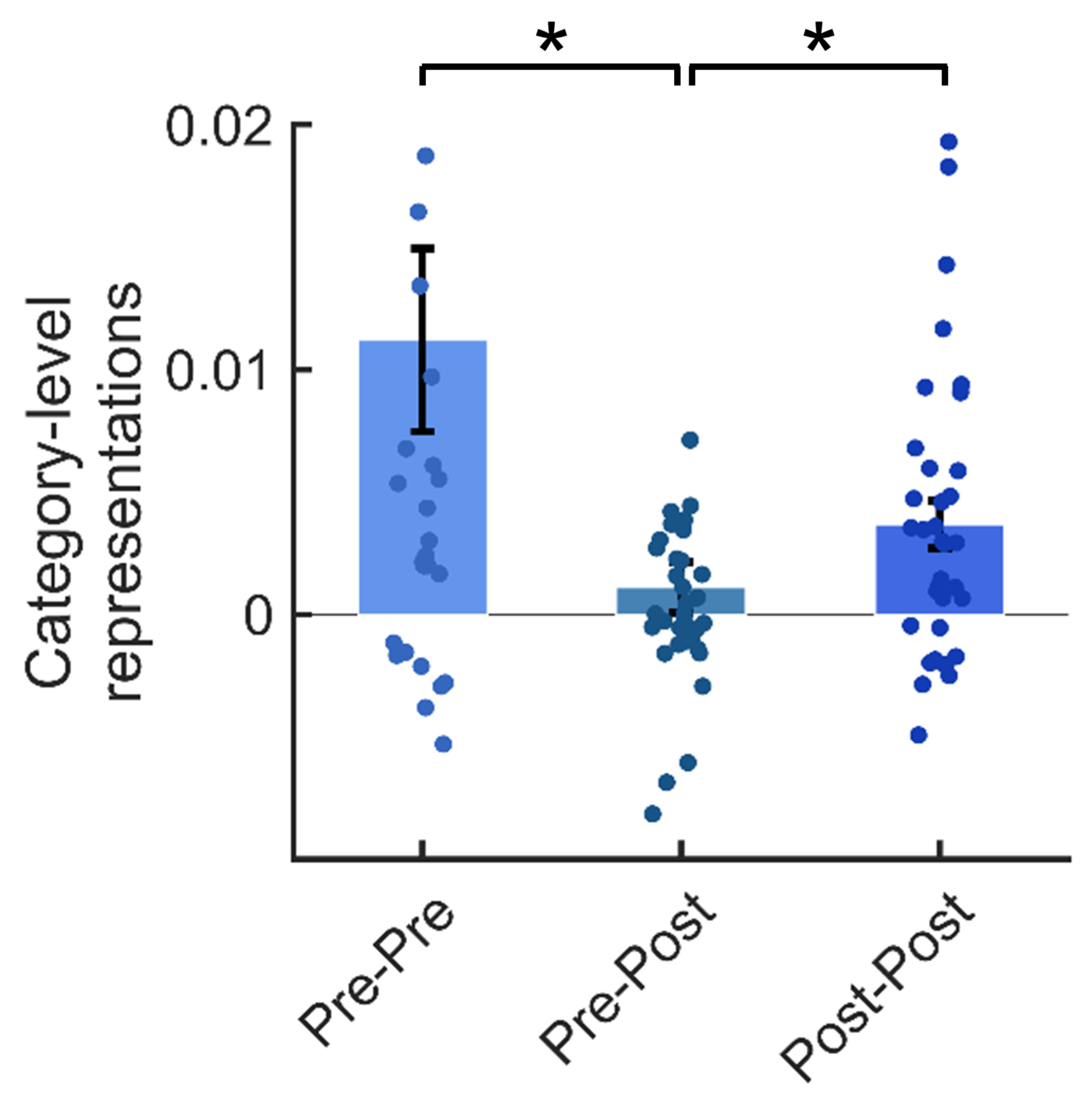

**Fig S3. Within-session category-level representations were greater than the cross-session category-level representation.** Providing significant category-level representational clusters were identified within both the pre-sleep learning and post-sleep mental retrieval sessions, we also contrasted the within-session category-level representations versus between-session category-level representations by recruiting the time windows of the identified clusters. Specifically, the category-level Pre-Pre similarity was obtained by averaging the similarity values within the cluster identified during the pre-sleep learning session (0-1000 ms and 3000-4800 ms post encoding onset, see Fig. 2B). Similarly, the category-level Post-Post similarity was obtained by averaging the similarity values within the cluster identified during the post-sleep retrieval session (900-2900 ms post retrieval cue onset, see Fig. 2F). The category-level Pre-Post similarity was determined by correlating the EEG power in the time windows of the pre-sleep learning cluster with that in the time windows of the post-sleep retrieval cluster. Results show that category-level Pre-Pre was significantly greater than the category-level Pre-Post similarity (*t*(34) = 2.60, *p* = 0.014), and Post-Post similarity was greater than Pre-Post similarity (*t*(34) = 2.22, *p* = 0.033).

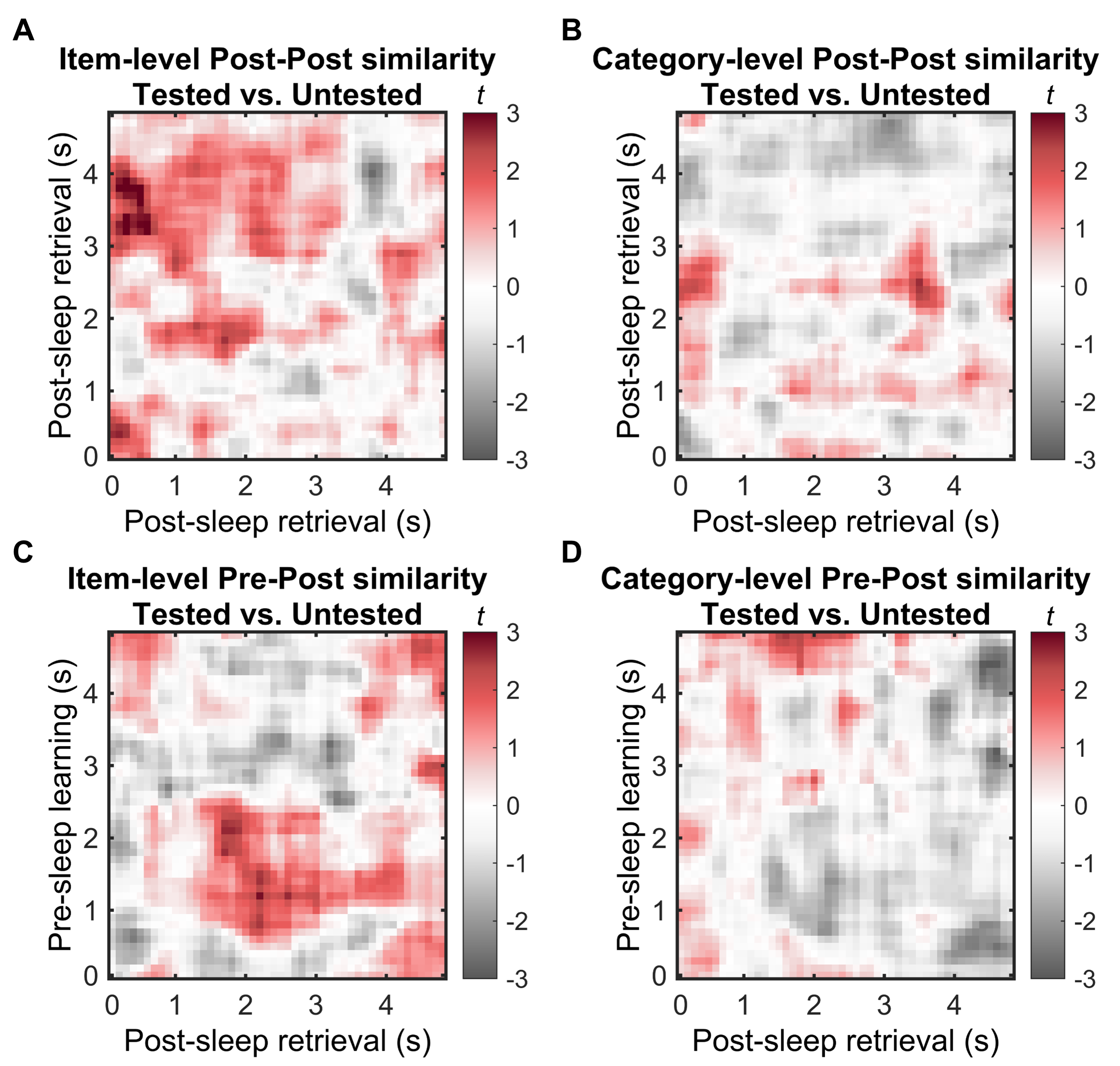

**Fig S4. Neural representations for pre-sleep tested versus untested items among post-sleep remembered items.** To understand whether pre-sleep testing affects memory representational transformation, we compared the neural representations for items that were tested pre-sleep (i.e., tested items) versus items that were not tested pre-sleep (i.e., untested items). Nine participants with fewer than ten trial pairs in the RSA for the post-sleep remembered untested condition were excluded, resulting in 26 participants in this analysis. **(A-B)** Within the post-sleep mental retrieval session (Post-Post similarity), there were no significant differences between tested and untested items for either the item-level (*p*_cluster_ > 0.152) or the category-level representations (*p*_cluster_ > 0.515). **(C-D)** For cross-session similarity (Pre-Post similarity), no significant differences were found between tested and untested items for either the item-level (*p*_cluster_ > 0.340) or the category-level representations (*p*_cluster_ > 0.394).

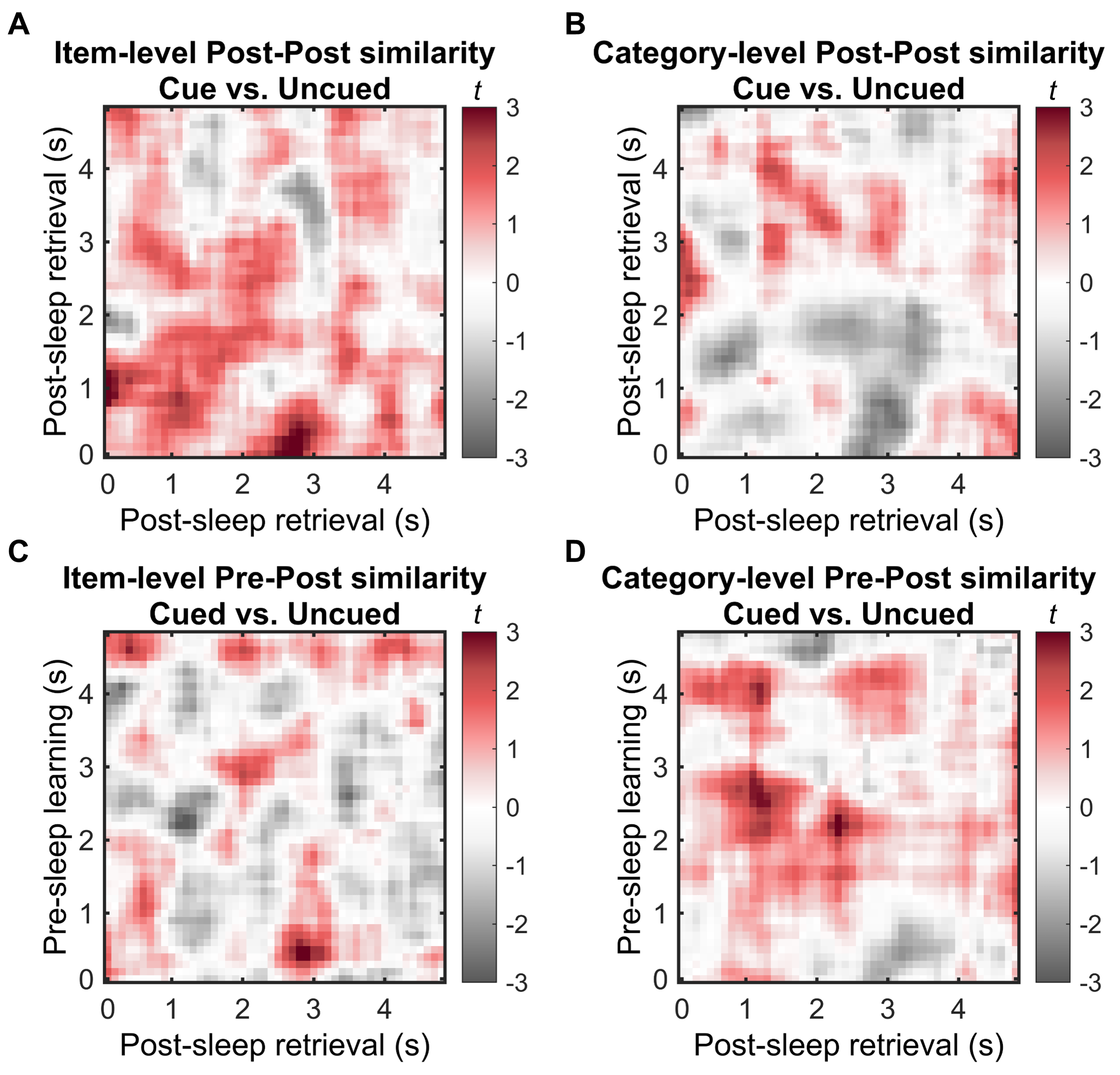

**Fig S5. Neural representations for TMR cued versus uncued items among post-sleep remembered items.** To understand whether targeted memory reactivation (TMR) during slow-wave sleep affects memory representational transformation, we compared neural representations for items cued during the TMR session versus uncued items. **(A-B)** Within the post-sleep mental retrieval session (Post-Post similarity), there were no significant differences between cued and uncued items for either the item-level (*p*_cluster_ > 0.196) or the category-level representations (*p*_cluster_ > 0.266). **(C-D)** For cross-session similarity (Pre-Post similarity), no significant differences were found between cued and uncued items for either the item-level (*p*_cluster_ > 0.469) or the category-level representations (*p*_cluster_ > 0.195).

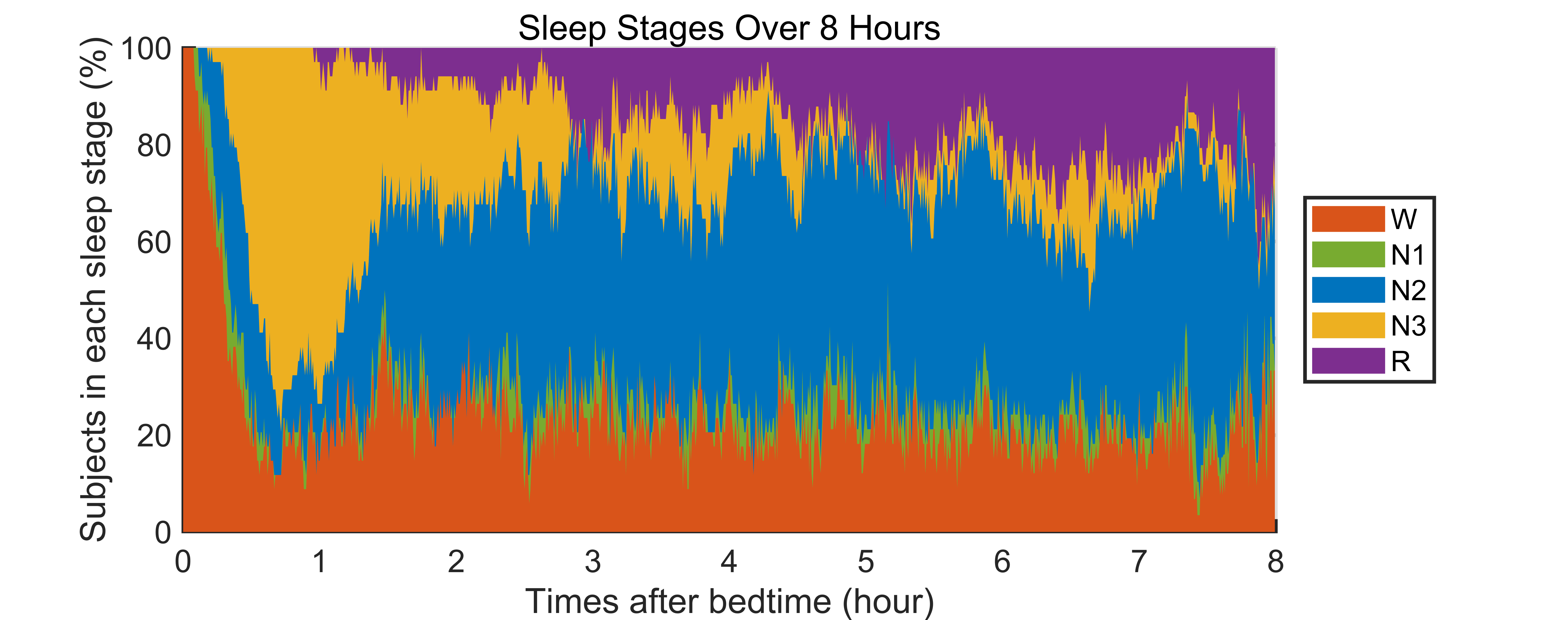

**Fig S6. Distribution of sleep stages over an 8-hour period.** W: wake, R: REM sleep.

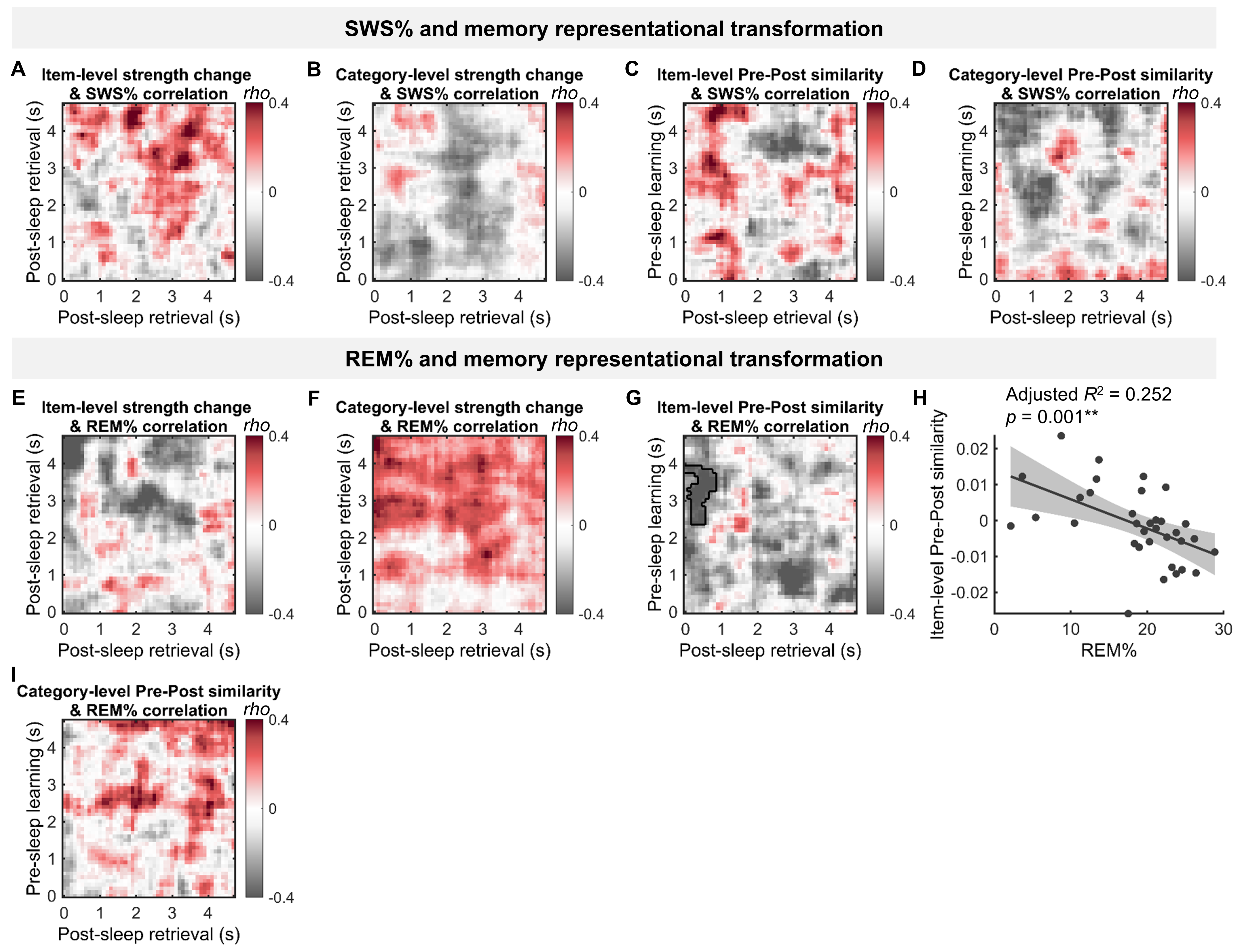

**Fig S7. The relationship between memory representational transformation and individual sleep stages: slow-wave sleep (SWS) and REM sleep. (A-B)** SWS% showed no significant correlation with either item-level or category-level representational strength change (Post minus Pre) (*p*s_cluster_ > 0.225). **(C-D)** SWS% showed no significant correlation with item-level or category-level Pre-Post representational similarity (*p*s_cluster_ > 0.198). **(E-F)** REM% showed no significant correlation with either item-level or category-level representational strength change (*p*_cluster_ > 0.075). **(G-H)** REM% was negatively correlated with item-level Pre-Post representational similarity (*p*_cluster_ = 0.04; within the cluster: *β* = -0.08, adjusted *R*^2^ = 0.252, *p* = 0.001). **(I)** REM% showed no significant correlation with category-level Pre-Post representational similarity (*p*_cluster_ > 0.570). Significant clusters with *p*_cluster_ < 0.05 were circled by black lines. **: *p* < 0.01.

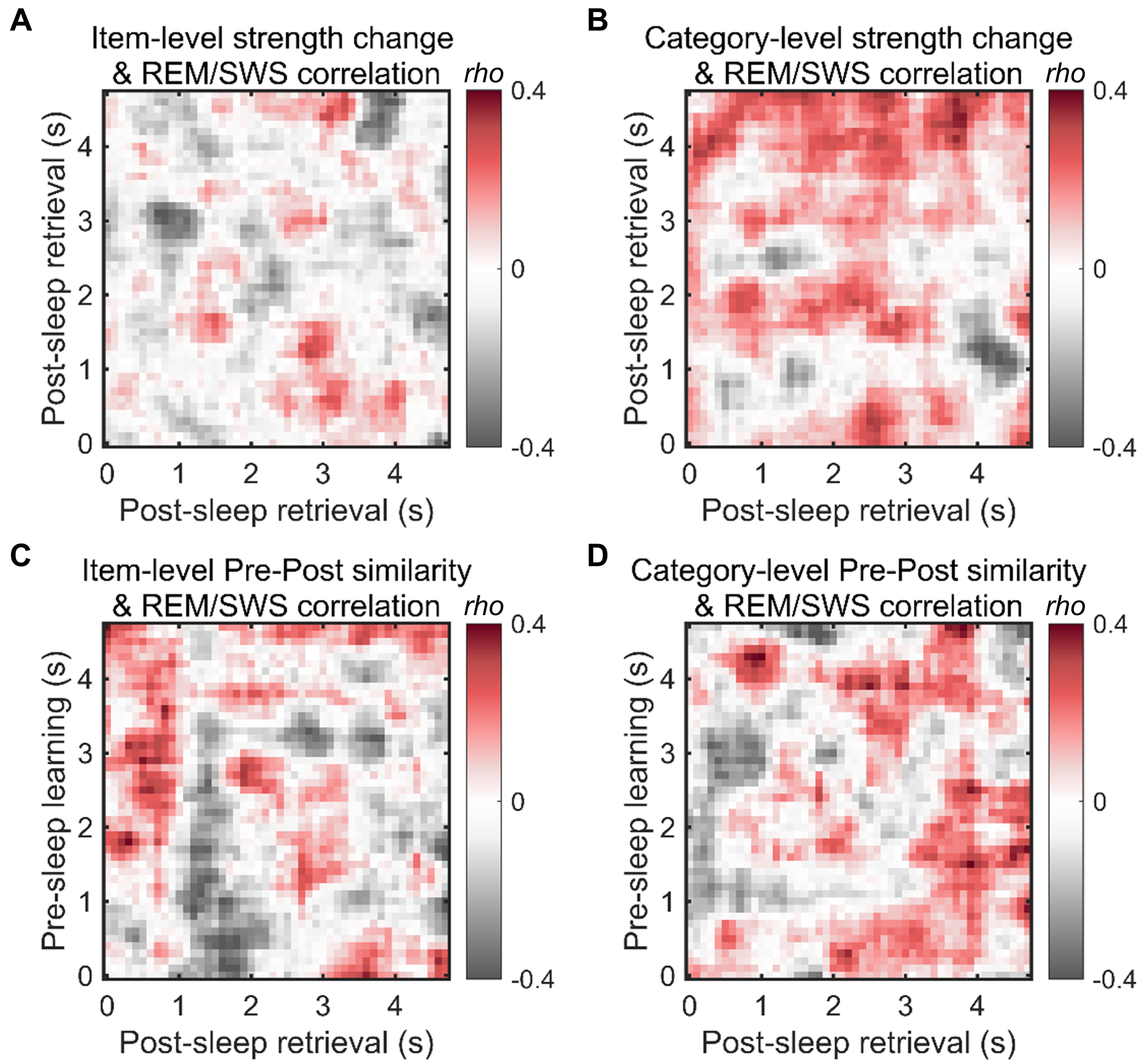

**Fig S8. REM/SWS showed no significant correlation with memory representational transformation indexes for post-sleep forgotten items.** No significant clusters were identified between REM/SWS and the following memory representational transformation indexes for post-sleep forgotten items: **(A)** item-level strength change (*p*_cluster_ > 0.534); **(B)** category-level strength change (*p*_cluster_ > 0.282); **(C)** item-level Pre-Post similarity (*p*_cluster_ > 0.473); **(D)** category-level Pre-Post similarity (*p*_cluster_ > 0.705).

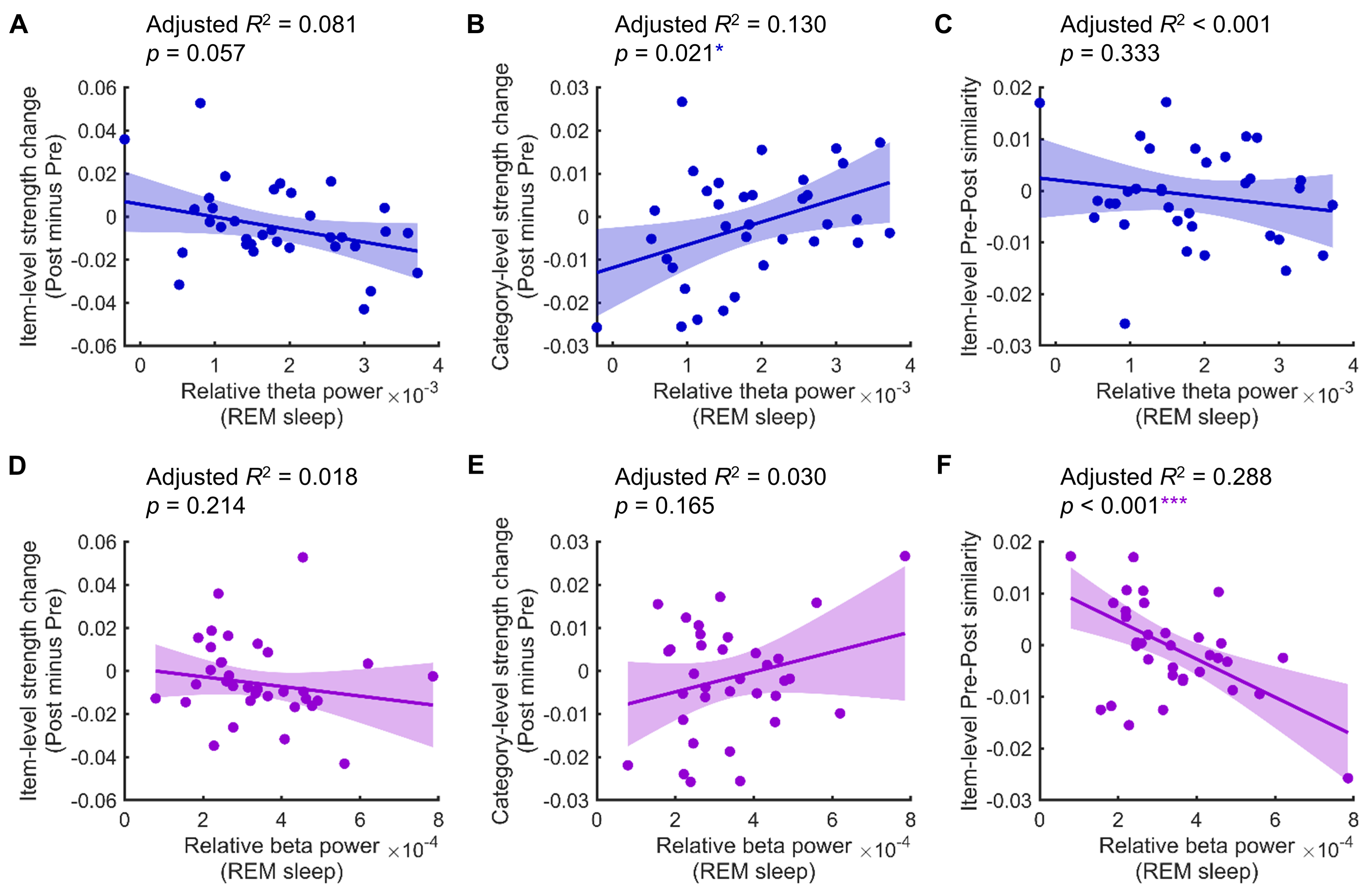

**Fig S9. Relationship between relative REM sleep power and memory representational transformation.** **(A-B)** Relative theta band (4-7 Hz) power from frontal electrodes (F3 and F4) during REM sleep showed a negative but non-significant correlation with item-level strength change (post minus pre), while showing a significant positive correlation with category-level strength change (post minus pre). **(C)** Relative theta power during REM sleep showed no significant correlation with item-level Pre-Post similarity. **(D-E)** Relative beta band (15-25 Hz) power from the same frontal electrodes during REM sleep showed no significant correlation with either item-level strength change or category-level strength change. **(F)** Relative theta power during REM sleep showed a significant negative correlation with item-level Pre-Post similarity.

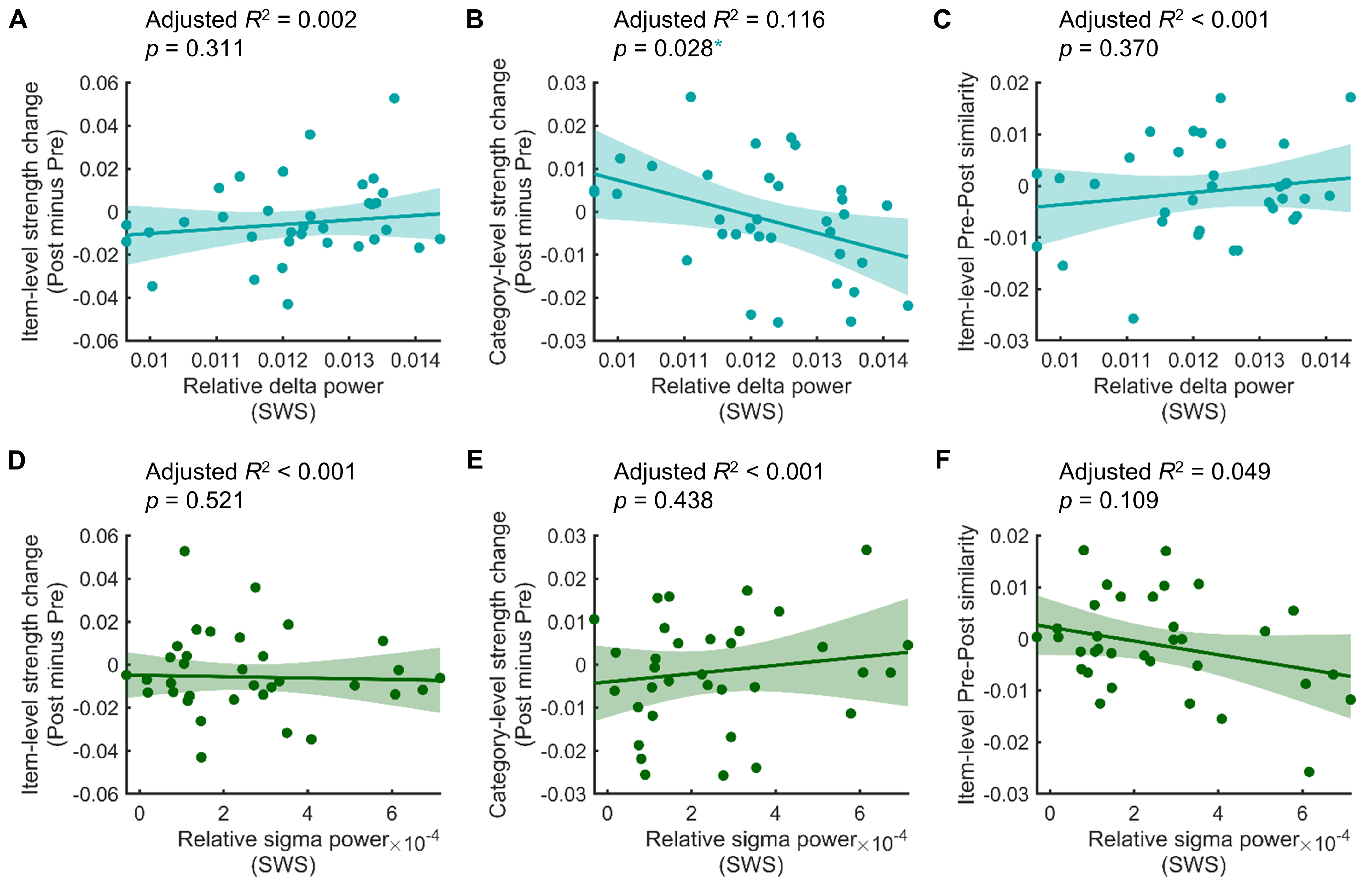

**Fig S10. Relationship between relative SWS power and memory representational transformation.** **(A-B)** Relative delta band (4-7 Hz) power from Fz and Cz electrodes during SWS showed no significant correlation with item-level strength change (post minus pre), but showed a significant negative correlation with category-level strength change (post minus pre). **(C)** Relative delta power during SWS showed no significant correlation with item-level Pre-Post similarity. **(D-F)** Relative sigma band (11-16 Hz) power from the same electrodes (Fz and Cz) during SWS showed no significant correlation with item-level strength change, category-level strength change, or item-level Pre-Post similarity.

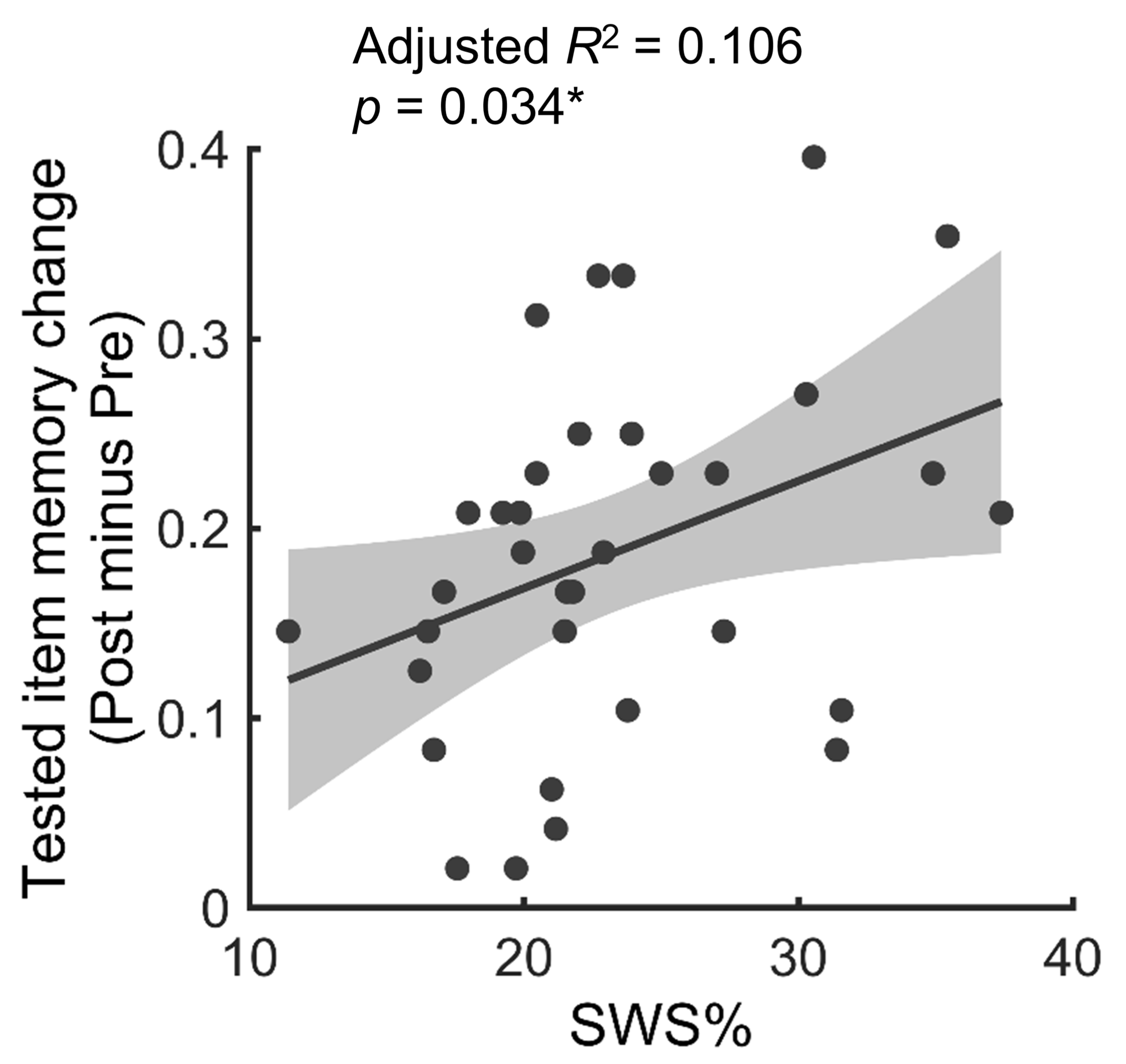

**Fig S11. Slow-wave sleep duration (%) was positively correlated with the memory performance change (post-sleep test minus pre-sleep test) for pre-sleep tested item.**

**Table S1.** **Sleep stage descriptive statistics** (*n* = 34)

| Sleep Statistics | Mean | SD |
| --- | --- | --- |
| Total time in bed (min) | 476.96 | 48.70 |
| Sleep duration (min) | 359.99 | 81.37 |
| Stage N1 duration (min) | 21.81 | 13.14 |
| Stage N2 duration (min) | 187.06 | 49.37 |
| Stage N3/SWS duration (min) | 82.13 | 22.98 |
| REM sleep duration (min) | 68.99 | 31.10 |
| Stage N1 (%) | 6.23 | 3.94 |
| Stage N2 (%) | 52.04 | 8.58 |
| Stage N3 (%) | 23.23 | 6.08 |
| REM sleep (%) | 18.50 | 6.65 |
